# To integrate or not to integrate: Temporal dynamics of hierarchical Bayesian Causal Inference

**DOI:** 10.1101/504118

**Authors:** Máté Aller, Uta Noppeney

**Author notes:** Corresponding author: Máté Aller, Computational Neuroscience and Cognitive Robotics Centre, University of Birmingham, Edgbaston, Birmingham B15 2TT, United Kingdom.

## Abstract

To form a percept of the environment, the brain needs to solve the binding problem – inferring whether signals come from a common cause and be integrated, or come from independent causes and be segregated. Behaviourally, humans solve this problem near-optimally as predicted by Bayesian Causal Inference; but, the neural mechanisms remain unclear. Combining Bayesian modelling, electroencephalography (EEG), and multivariate decoding in an audiovisual spatial localization task, we show that the brain accomplishes Bayesian Causal Inference by dynamically encoding multiple spatial estimates. Initially, auditory and visual signal locations are estimated independently; next, an estimate is formed that combines information from vision and audition. Yet, it is only from 200 ms onwards that the brain integrates audiovisual signals weighted by their bottom-up sensory reliabilities and top-down task-relevance into spatial priority maps that guide behavioural responses. Critically, as predicted by Bayesian Causal Inference, these spatial priority maps take into account the brain’s uncertainty about the world’s causal structure and flexibly arbitrate between sensory integration and segregation. The dynamic evolution of perceptual estimates thus reflects the hierarchical nature of Bayesian Causal Inference, a statistical computation, crucial for effective interactions with the environment.

## Introduction

In our natural environment our senses are exposed to a barrage of sensory signals: the sight of a rapidly approaching truck, its looming motor noise, the smell of traffic fumes. How the brain effortlessly merges these signals into a seamless percept of the environment remains unclear. Critically, the brain faces two fundamental computational challenges: First, we need to solve the ‘binding’ or ‘causal inference’ problem - deciding whether signals come from a common cause and thus should be integrated, or instead be treated independently [1,2]. Second, when there is a common cause, the brain should integrate signals taking into account their uncertainties [3,4]).

Hierarchical Bayesian Causal Inference provides a rational strategy to arbitrate between sensory integration and segregation in perception [2]. Bayesian Causal Inference explicitly models the potential causal structures that could have generated the sensory signals i.e., whether signals come from common or independent sources. In line with Helmholtz’s notion of ‘unconscious inference’, the brain is then thought to invert this generative model during perception [5]. In case of a common signal source, signals are integrated weighted in proportion to their relative sensory reliabilities (i.e. forced fusion [3,4,6–10]). In case of independent sources, they are processed independently (i.e. full segregation [11,12]). Critically, on a particular instance the brain does not know the world’s causal structure that gave rise to the sensory signals. To account for this causal uncertainty a final estimate (e.g. object’s location) is obtained by averaging the estimates under the two causal structures (i.e. common vs. independent source models) weighted by each causal structure’s posterior probability – a strategy referred to as model averaging (for other decisional strategies see [13]).

A large body of psychophysics research has demonstrated that human observers combine sensory signals near-optimally as predicted by Bayesian Causal Inference [2,13–16]. Most prominently, when locating events in the environment observers gracefully transition between sensory integration and segregation as a function of audiovisual spatial disparity [12]. For small spatial disparities, they integrate signals weighted by their reliabilities leading to crossmodal spatial biases [17], for larger spatial disparities audiovisual interactions are attenuated. A recent fMRI study showed how Bayesian Causal Inference is accomplished within the cortical hierarchy [14,16]: While early auditory and visual areas represented the signals on the basis that they were generated by independent sources (i.e. full segregation), the posterior parietal cortex integrated sensory signals into one unified percept (i.e. forced fusion). Only at the top of the cortical hierarchy, in anterior parietal cortex, the uncertainty about the world’s causal structure was taken into account and signals were integrated into a spatial estimate consistent with Bayesian Causal Inference.

The organization of Bayesian Causal Inference across the cortical hierarchy raises the critical question of how these neural computations unfold dynamically over time within a trial. How does the brain merge spatial information that is initially coded in different reference frames and representational formats? While the brain is likely to recurrently update all spatial estimates by passing messages forwards and backwards across the cortical hierarchy [18–20], the unisensory estimates may to some extent precede the computation of the Bayesian Causal Inference estimate. To characterize the neural dynamics of Bayesian Causal Inference we presented human observers with auditory, visual and audiovisual signals that varied in their spatial disparity in an auditory and visual spatial localization task whilst recording their neural activity with EEG. First, we employed cross-sensory decoding and temporal generalization matrices [21] of the unisensory auditory and visual signal trials to characterize the emergence and the temporal stability of spatial representations across the senses. Second, combining psychophysics, EEG and Bayesian modelling, we temporally resolved the evolution of unisensory segregation, forced fusion and Bayesian causal inference in multisensory perception.

## Results

To determine the computational principles that govern multisensory perception we presented 13 participants with synchronous audiovisual spatial signals (i.e. white noise burst and Gaussian cloud of dots) that varied in their audiovisual spatial disparity and visual reliability (Fig. 1A, B). On each trial, participants reported their perceived location of either the auditory or the visual signal. In addition, we included unisensory auditory and visual signal trials under auditory or visual report, respectively.

Combining psychophysics, EEG and computational modelling, we addressed two questions: First, we investigated when and how human observers form spatial representations from unisensory visual or auditory inputs, which generalize across the two sensory modalities. Second, we studied the computational principles and neural dynamics that mediate the integration of audiovisual signals into spatial representations that take into account the observer’s uncertainty about the world’s causal structure consistent with Bayesian Causal Inference.

**Figure 1.**
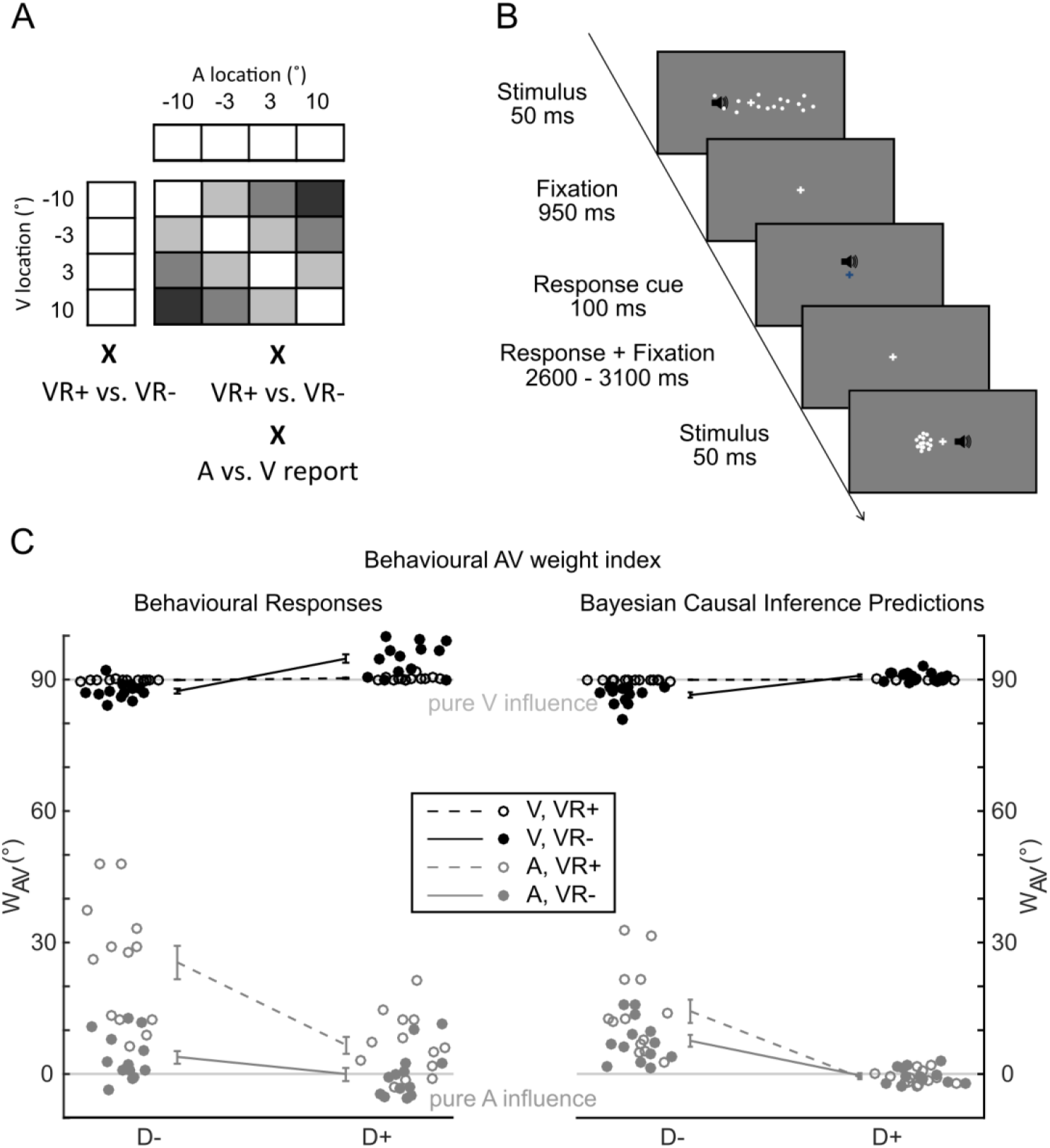
Experimental design, example trial, behavioural and predicted audiovisual weights (w_AV_) (**A**) Experimental design. In a 4 x 4 x 2 x 2 factorial design, the experiment manipulated i. the location of the visual (V) signal (-10°, -3.3°, 3.3°, and 10°), ii. the location of the auditory (A) signal (-10°, -3.3°, 3.3°, and 10°), iii. the reliability of the visual signal (high [VR+] versus low [VR-], as defined by the spread of the visual cloud), and iv. task relevance (auditory versus visual report). In addition, we included unisensory auditory and visual [VR+] and [VR-] trials. The greyscale codes the spatial disparity between the auditory and visual locations for each AV condition (i.e. darker greyscale = larger spatial disparity). (**B**) Time course of an example trial. (**C**) Behavioural audiovisual weight index w_AV_ computed from behavioural responses (left) and from the predictions of the Bayesian Causal Inference model (right; across-participants circular mean ± 68% CI and individual w_AV_ represented by filled/empty circles, n = 13). The audiovisual weight index wAV is shown as a function of (i) visual reliability: high [VR+] versus low [VR-], (ii) task relevance: auditory [A] versus visual [V] report, (iii) audiovisual spatial disparity: small [≦6.6; D-] versus large [>6.6; D+].

### Shared and distinct neural representations of space across vision and audition – unisensory auditory and visual conditions

#### Behavioural results

Participants were able to locate unisensory auditory and visual signals reliably as indicated by a significant Pearson correlation between participant’s location responses and the true signal source location for both unisensory auditory (across subjects mean ± SEM: 0.88 ± 0.05), visual high reliability [VR+] (across subjects mean ± SEM: 0.998 ± 0.19) and visual low reliability [VR-] (across subjects mean ± SEM: 0.91 ± 0.05) conditions. As expected, observers were significantly less accurate when locating the sound than when locating the visual stimuli for both levels of visual reliability (VR+ vs A: t(12) 8.83, p < 0.0001; VR-vs. A: t(12) = 1.47, p = 0.005; see Fig. S1 for response distributions across all conditions).

#### EEG results

Multivariate decoding of EEG activity patterns revealed how the brain dynamically encodes the location of unisensory auditory or visual signals. The decoding accuracy was expressed as the Pearson correlation coefficient between the true and the decoded stimulus locations and entered into so-called temporal generalization matrices that illustrate the stability of EEG activity patterns encoding spatial location across time [21]. If a SVR model trained on EEG activity patterns at time t can correctly decode the stimulus location not only at time t but also at other time points, then the stimulus location is encoded in EEG activity patterns that are relatively stable across time (for further details about this temporal generalization approach see [21]). If an SVR model cannot successfully generalize to EEG activity patterns at other time points, spatial locations are encoded in transient EEG activity patterns that differ across time.

For visual stimuli, spatial locations were successfully (i.e. significantly better than chance) decoded from EEG activity patterns from 60 ms onwards for visual stimuli (Fig 2, upper right quadrant and Suppl. Fig S2A). Moreover, the temporal generalization matrices suggest that the visual spatial representations were initially transient (i.e. 60 - 150 ms, significant decoding accuracy only near the diagonal) reflecting the early visual evoked EEG responses (e.g. N1, see Fig S2A, B). Later (i.e. from about 250 ms) the location of the visual stimulus was encoded in a more sustained activity pattern (see Fig S2A, B) leading to successful cross-temporal generalization from 300 ms to 700 ms poststimulus (i.e. significantly better than chance decoding accuracy is present far off the diagonal).

By contrast, auditory spatial representations could be decoded significantly better than chance from about 95 ms onwards (see Fig 2, lower left quadrant), which corresponds to the auditory N1 component (Fig S3A, B). Particularly from 200 ms onwards the SVR decoder trained on EEG activity patterns can decode auditory spatial location significantly better than chance also from EEG activity patterns across other time points, even as late as 700 ms post stimulus (significant cluster encircled by thin grey lines). This temporal generalization profile indicates that auditory spatial locations were encoded in EEG activity patterns that were relatively stable across time later from 200 ms onwards. Visual inspection of the EEG topographies shows that auditory spatial location is encoded at these later processing stages in sustained activity patterns that correspond to the long latency auditory P2 component (see Fig S3A, B) [22–24].

**Figure 2.**
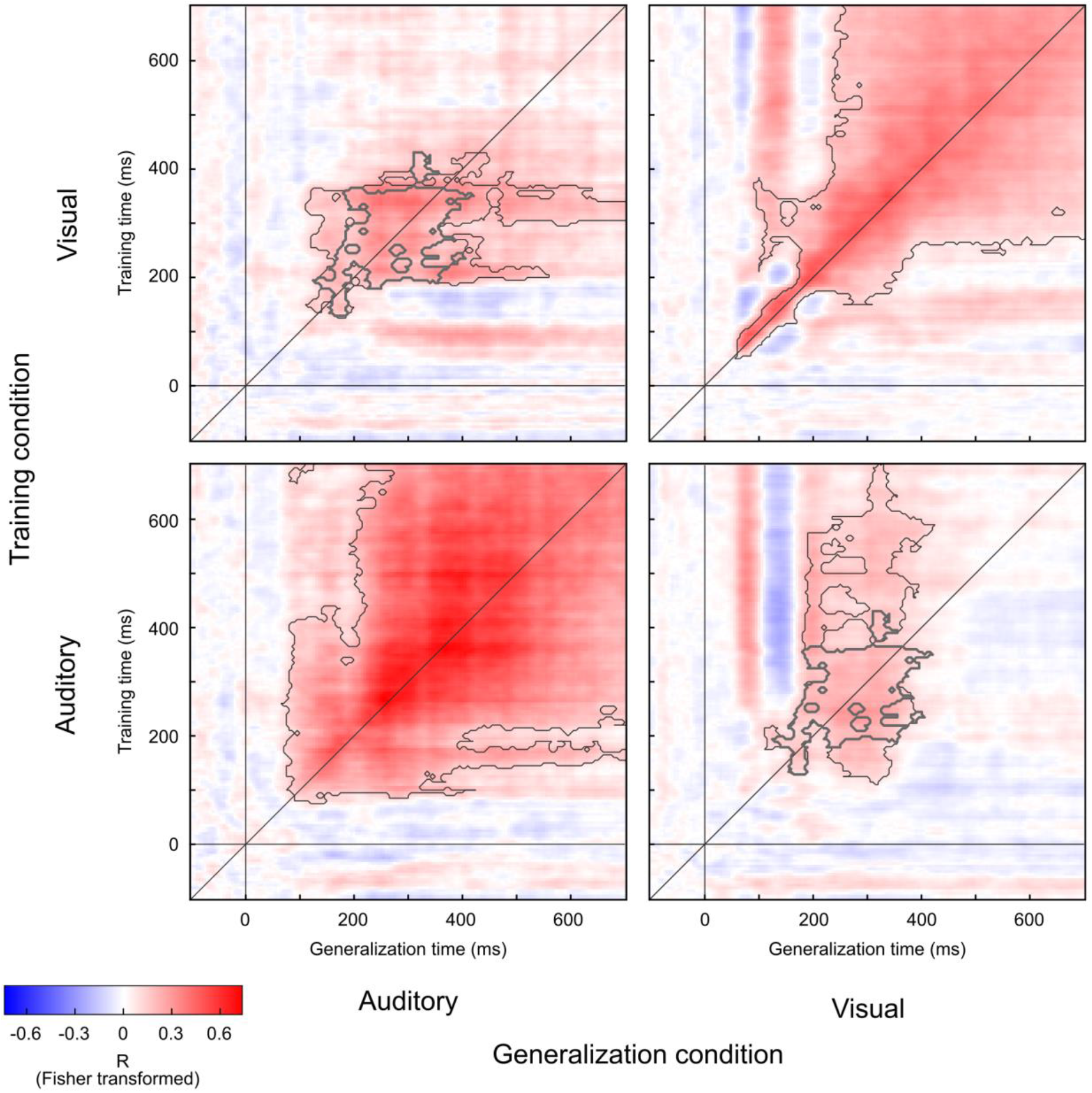
Temporal generalization matrices within and across auditory and visual senses. Each temporal generalization matrix shows the decoding accuracy for each training (y axis) and testing (x axis) time point. We factorially manipulated the training data (auditory vs. visual stimulation) and testing data (auditory vs. visual stimulation). Decoding accuracy is quantified by the Pearson correlation between the true and the decoded locations of the auditory (resp. visual) stimulus. The grey line along the diagonal indicates where the training time is equal to the testing time (i.e. the time resolved decoding accuracies). Horizontal and vertical grey lines indicate the stimulus onset. The thin grey lines encircle clusters with decoding accuracies that were significantly better than chance at p < 0.05 corrected for multiple comparisons. The thick grey lines encircle the clusters with decoding accuracies that were significantly better than chance jointly for both i. auditory to visual and ii. visual to auditory cross-temporal generalization at p < 0.05 corrected for multiple comparisons.

Critically, in addition to temporal generalization within each sensory modality, we also investigated the extent to which the SVR decoding model generalized across sensory modalities throughout post-stimulus time. While earlier neural representations were more specific to each particular sensory modality, the SVR model was able to generalize significantly better than chance from audition to vision and vice versa from 160 to 360 ms (Fig 2, upper left and lower right quadrant, areas encircled by thick grey line indicate significant generalization across sensory modalities). This cross-sensory generalization across visual and auditory-evoked EEG activity patterns suggests that at those stages (i.e. 160 ms to 360 ms) the brain forms spatial representations that are relatively stable and rely on neural generators that may be partly shared across sensory modalities. By contrast, the spatial representations encoded in very early (<160 ms) EEG activity patterns did not enable successful cross-sensory generalization suggesting that they are modality-specific. These statistically significant cross-sensory generalization results are also illustrated by the EEG topographies evoked by unisensory auditory and visual signals (cf. Fig S2B, S3B). From 200 ms to 400 ms post-stimulus auditory and visual stimuli elicit centro-posterior dominant topographies that depend on the stimulus location to some extent similarly in vision and audition. While these results may point towards partly overlapping neural generators and representations potentially in parietal cortices that encode location both in audition and vision, it is important to emphasize that different configurations of neural generators can in principle elicit similar EEG scalp topographies.

### Computational principles of audiovisual integration: GLM-based w_AV_ and Bayesian modelling analysis – audiovisual conditions

Combining psychophysics, multivariate EEG pattern decoding and computational modelling, we next investigated the computational principles and neural dynamics underlying audiovisual integration of spatial representations using a general linear model (GLM)-based w_AV_ and a Bayesian modelling analysis. Critically, as shown in Fig 3 both analyses were applied to the spatial estimates that were either reported by participants (i.e. behaviour, Fig 3B left) or decoded from EEG activity patterns independently for each post-stimulus time point (i.e. neural, Fig 3B right, for further details see methods section and figure legend).

**Figure 3.**
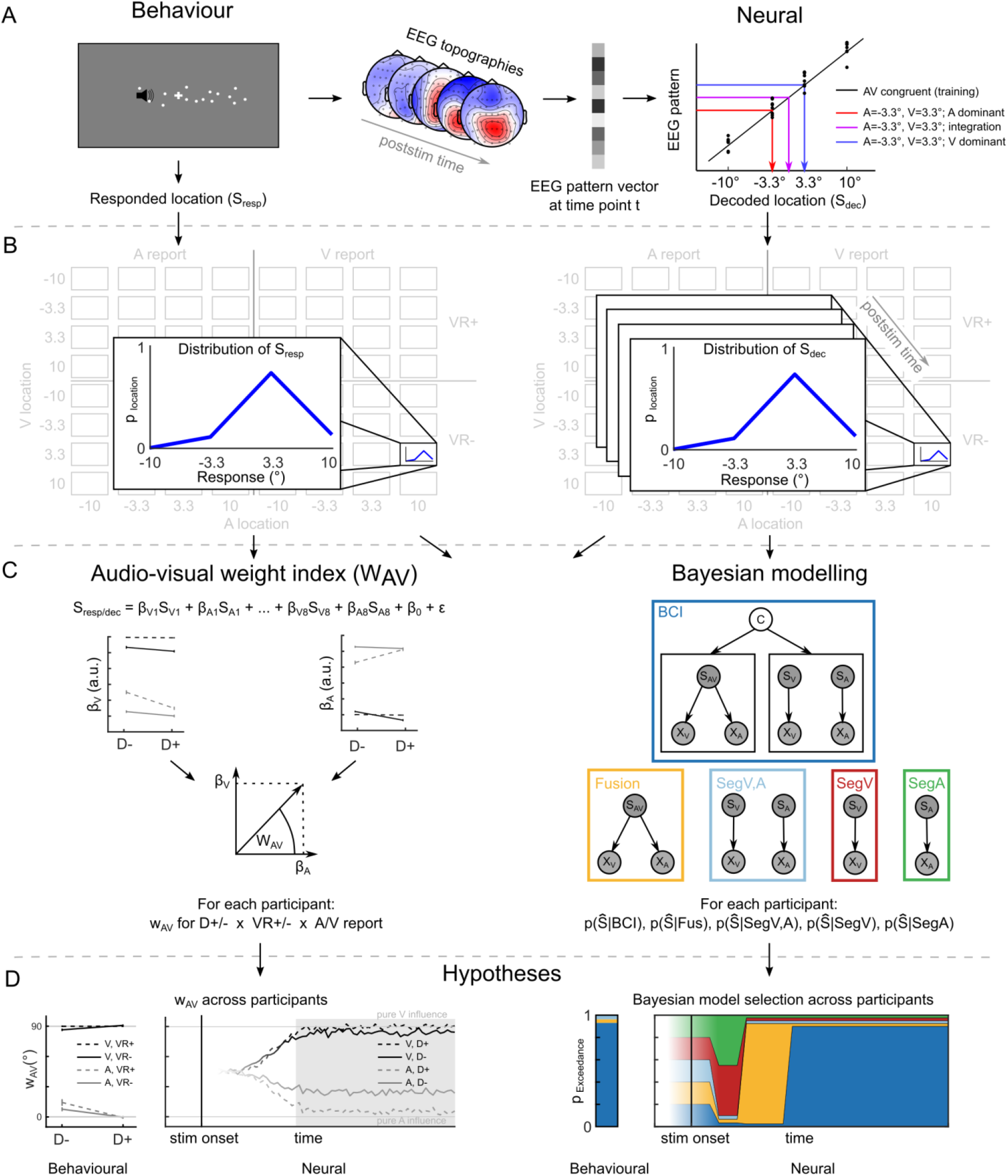
GLM-based w_AV_ and Bayesian modelling analysis overview. (**A**) The GLM-based w_AV_ and Bayesian modelling analysis were performed on auditory and visual spatial estimates that were indicated by participants as behavioural localization responses (left, Behaviour) or decoded from participants’ EEG activity patterns (right, Neural). The neural spatial estimates were obtained by training a SVR model on ERP activity patterns at each time point of the audiovisual congruent trials to learn the mapping from EEG pattern to external spatial locations (black diagonal line). This learnt mapping was then used to decode the spatial location from the ERP activity patterns of the spatially congruent and incongruent audiovisual conditions (coloured arrows). (**B**) Distributions of spatial localization responses (left, Behaviour: S_Resp_= Spatial estimate responded) and decoded spatial estimates (right, Neural: S_Dec_= Spatial estimate decoded) were computed for each of the 64 conditions of the the 4 (visual stimulus location) x 4 (auditory stimulus location) x 2 (visual reliability) x 2 (task relevance) factorial design. (**C left**) In the GLM-based w_AV_ analysis the perceived (or decoded at each time point) spatial estimates were predicted by the true visual and auditory spatial locations (S_V1..8_, S_A1..8_) for each of the eight conditions in the 2 (visual reliability: high vs. low) x 2 (task-relevance: auditory vs. visual report) x 2 (spatial disparity: < 6.6° vs. ≤ 6.6°) factorial design. As a summary index we defined the relative audiovisual weight (w_AV_) as the four-quadrant inverse tangent of the visual (ß_V1..8_) and auditory (ß_A1..8_) parameter estimates for each of the eight conditions in each regression model. (**C right**) In the Bayesian-modelling analysis we fitted the following models to observers’ behavioural and neural responses: ‘segregation unisensory auditory’ (SegA, green, for EEG only), ‘segregation unisensory visual’ (SegV, red, for EEG only), ‘full audiovisual segregation’ (SegV,A, light blue), ‘forced fusion’ (Fusion, yellow) and ‘Bayesian Causal Inference’ model (with model averaging, BCI, dark blue). We performed Bayesian model selection at the group level and computed the protected exceedance probability that one model is better than any of the other candidate models above and beyond chance [76]. (**D left**) Based on previous studies ([14,16]) we hypothesized that the qualitative w_AV_ profile with an interaction between task-relevance (i.e. visual vs. auditory report) and spatial disparity that is characteristic for Bayesian Causal Inference would emerge relatively late. (**D right**) Likewise, we expected the different models to dominate the EEG activity patterns to some extent sequentially: first the unisensory segregation model (SegV,SegA), followed by the forced fusion model (Fusion) and finally the Bayesian Causal Inference estimate (BCI). The fading of colours indicates that we did not have specific hypotheses for those times.

The GLM-based w_AV_ analysis quantifies the influence of the true auditory and true visual location on (i) the reported or (ii) EEG decoded auditory and visual spatial estimates in terms of an audiovisual weight index w_AV_.

The Bayesian modelling analysis formally assessed the extent to which (i) the full segregation model(s) (Fig 3C, encircled in light blue, red or green) (ii) the forced fusion model (Fig 3C, yellow) and (iii) the Bayesian Causal Inference model (i.e., using model averaging as decision function, encircled in dark blue; see supporting material Table S1 for other decision functions) can account for the spatial estimates reported by observers (i.e. behavior) or decoded from EEG activity pattern (i.e. neural).

#### Behavioural results

##### GLM-based w_AV_ analysis

The behavioural audiovisual weight index w_AV_ shows that observers integrated audiovisual signals weighted by their sensory reliabilities and task relevance (see Fig 1C and Fig S1 for histograms of reported signal locations across all conditions).

The audiovisual weight index w_AV_ was close to 90° (i.e. pure visual influence), when the visual signal needed to be reported (Fig 1C, dark lines). But it shifted towards 0° when the auditory signal was task-relevant (Fig 1C, grey lines). In other words, we observed a significant main effect of task-relevance on behavioural w_AV_ (p = 0.0002). Observers flexibly adjusted the weights they assigned to auditory and visual signals in the integration process as a function of task-relevance giving more emphasis to the sensory modality that needed to be reported. The main effect of task-relevance on w_AV_ is inconsistent with classical forced fusion models where audiovisual signals are integrated into one single unified percept irrespective of task-relevance of the sensory modalities. In other words, even in the case of audiovisual spatial disparity, the observer would perceive the auditory and visual signals at the same location. Instead, it indicates that observers maintain separate auditory and visual spatial estimates for an audiovisual spatially disparate stimulus.

Critically, consistent with Bayesian Causal Inference the difference in w_AV_ between auditory and visual report significantly increased for large (> 6.6°) relative to small (≤ 6.6°) spatial disparities (i.e. significant interaction between task-relevance and spatial disparity: p = 0.0002). In other words, audiovisual integration and crossmodal spatial biasing broke down, when auditory and visual signals were far apart and likely to be caused by independent sources. This attenuation of audiovisual interactions for large relative to small spatial disparities (i.e. interaction between task-relevance and disparity) is the characteristic profile of Bayesian Causal inference (see model predictions for w_AV_ in Fig 1C right).

Moreover, we observed significant two-way interactions between visual reliability and spatial disparity (p = 0.0014) and between visual reliability and task-relevance (p = 0.0002). The effect of high vs. low visual reliability was stronger when the two signals were close in space and the auditory, i.e. less reliable signal needed to be reported. For auditory report conditions the influence of the visual signal on the audiovisual spatial representation is stronger for high visual reliability and small disparity trials (Fig 1C, difference between dashed and solid grey line for D-condition). Again, this interaction is expected for Bayesian Causal Inference, because the spatial estimate furnished by the forced fusion model receives a stronger weight in Bayesian Causal Inference for low spatial disparity trials, when it is likely that the two signals come from a common source.

##### Bayesian modeling analysis

Consistent with the profile of the audiovisual weight index w_AV_ formal Bayesian model comparison showed that the Bayesian Causal Inference model outperformed the full segregation and forced fusion models (85.6 ± 0.3 % variance explained, protected exceedance probability > 0.99; Table 1). Fig 1C (right) shows the profile of the audiovisual weight index w_AV_ that is predicted by the Bayesian Causal Inference model fitted to observer’s behavioural localization responses. It illustrates that Bayesian Causal Inference inherently accounts for effects of task-relevance (or reported modality) and the interaction between task-relevance and spatial disparity by combining the forced fusion estimate with the task-relevant full segregation estimate weighted by the posterior probability of common and independent sources. Conversely, the interaction between reliability and spatial disparity arises because the forced fusion model component which integrates signals weighted by their reliabilities is more dominant for small spatial disparities.

**Table 1.**
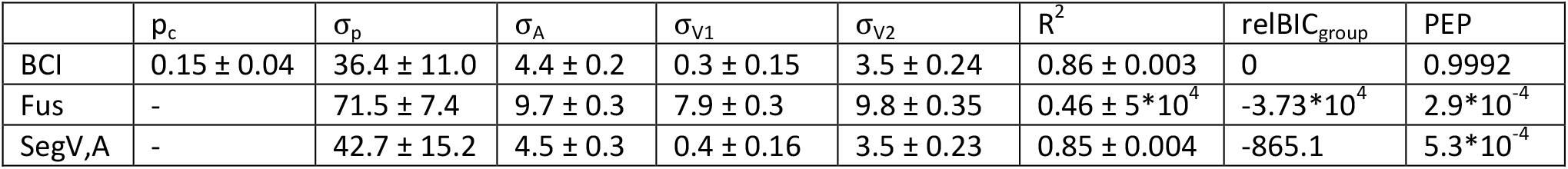
Model parameters (across-subjects mean ± SEM) of the computational models fit to observers’ behavioural localizations reports. R^2^ = coefficient of determination, relBIC_group_ = group level relative BIC, PEP = protected exceedance probability [76]. BCI = Bayesian Causal Inference Model; Fus = Fusion Model; SegV,A = Full segregation audiovisual model

In summary, our audiovisual weight index w_AV_ and Bayesian modelling analysis of observers’ perceived/reported locations provided convergent evidence that human observers integrate audiovisual spatial signals weighted by their relative reliabilities at small spatial disparities. Yet, they mostly segregate audiovisual signals at large spatial disparities, when it is unlikely that signals come from a common source.

#### EEG results - Temporal dynamics of audiovisual integration

To characterize the neural dynamics underlying integration of audiovisual signals into spatial representations, we applied the GLM-based wAV and the Bayesian modelling analysis to the ‘spatial estimates’ that were decoded from EEG activity patterns at each time point (see Fig 3B right). Because both the GLM-based wAV and the Bayesian modelling analysis require reliable spatial estimates, we report and interpret results limited to the time window from 55 ms to 700 ms post-stimulus (Fig 4, Fig S4), where the location of congruent audiovisual stimuli could be decoded better than chance from EEG activity patterns (p < 0.001).

**Figure 4.**
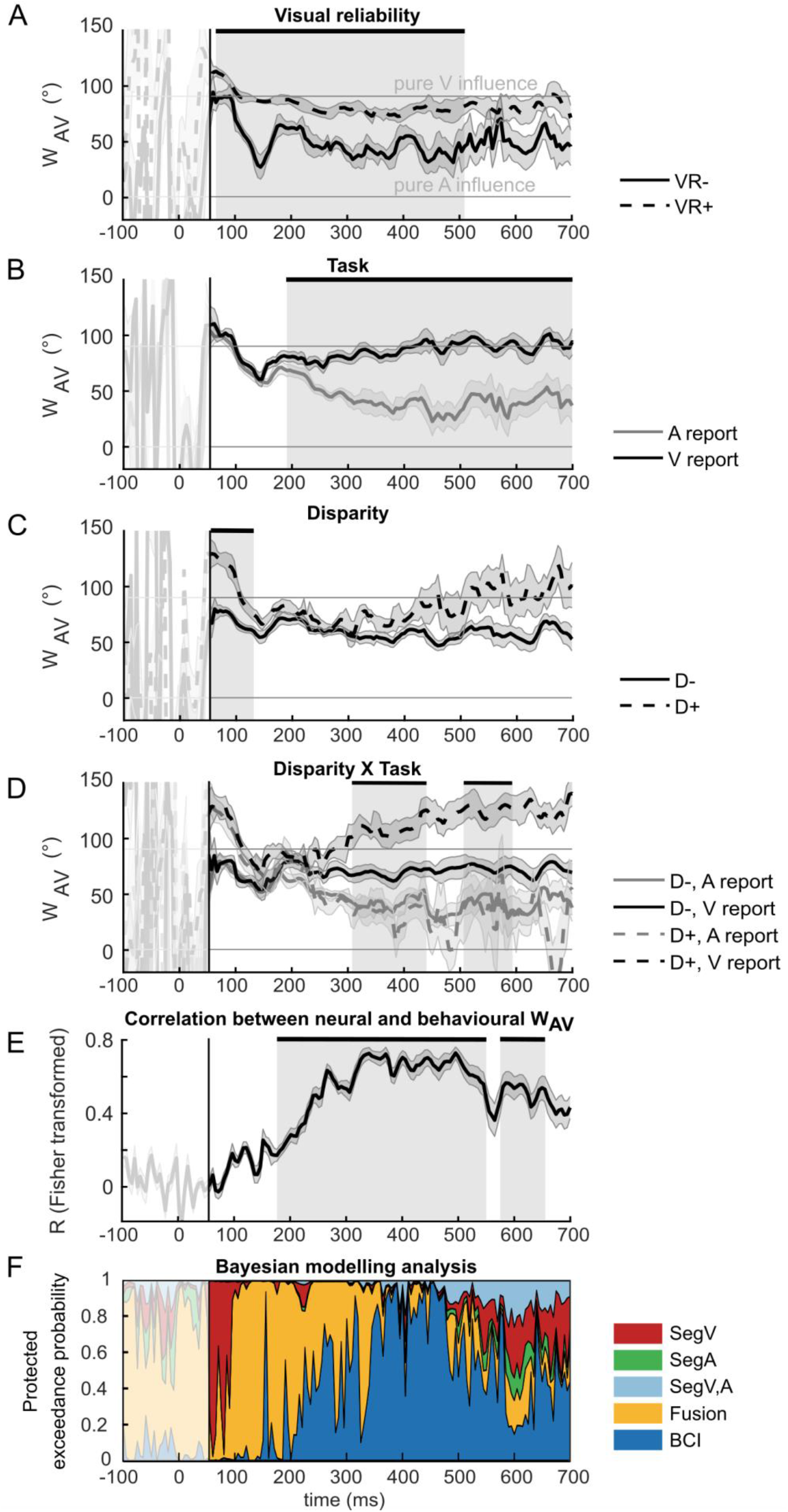
EEG results for GLM-based w_AV_ and Bayesian modelling analysis. The neural audiovisual weight index w_AV_ (across-participants’ circular mean ± 68% CI; n = 13). Neural w_AV_ as a function of time is shown for (**A**) visual reliability: high [VR+] versus low [VR-], (**B**) task relevance: auditory [A] versus visual [V] report, (**C**) audiovisual spatial disparity: small [≦6.6; D-] versus large [>6.6; D+], (**D**) the interaction between task relevance x disparity. Shaded grey areas indicate the time windows where the main effect of (**A**) visual reliability, (**B**) task relevance, (**C**) audiovisual spatial disparity or (**D**) the interaction between task relevance x disparity on w_AV_ was statistically significant at p < 0.05 corrected for multiple comparisons across time. (**E**) Time course of the circular-circular correlation (across-participants mean after Fisher z-transformation ± 68% CI; n = 13) between the neural- and the behavioural audiovisual weight index w_AV_. Shaded grey areas indicate significant correlation at p < 0.05 corrected for multiple comparisons across time. (**F**) Time course of the protected exceedance probabilities [76] of the five models of the Bayesian modelling analysis: ‘segregation unisensory auditory’ (SegA, green), ‘segregation unisensory visual’ (SegV, red), ‘full audiovisual segregation’ (SegV,A, light blue), ‘forced fusion’ (Fusion, yellow) and ‘Bayesian Causal Inference’ model (with model averaging, BCI, dark blue). The early time window until 55 ms (delimited by black vertical line on all plots) is shaded in white, because the decoding accuracy was not greater than chance for audiovisual congruent trials, hence the neural weight index w_AV_ and Bayesian model fits are not interpretable in this window.

##### GLM-based analysis of audiovisual weight index w_AV_

The GLM-based analysis investigated the effects of visual reliability, task-relevance and spatial disparity on the audiovisual neural weight index w_AV_ that quantifies the influence of auditory and visual signals on the spatial representations decoded from EEG activity patterns. Our results show that sensory reliability significantly influenced the neural w_AV_ from 65 – 510 ms. As expected, the spatial representations were more strongly influenced by the true visual signal location, when the visual signal was reliable than unreliable (i.e. significant main effect of visual reliability, Fig 4A, Table 2). Moreover, consistent with our behavioural findings, we also observed a significant main effect of task-relevance between 190 – 700 ms (Fig 4B, Table 2). As expected, the decoded location was more strongly influenced by the visual signal when the visual modality was task relevant. Most importantly, we also observed a significant interaction between task relevance and spatial disparity from 310 – 440 ms and 510 – 590 ms. As discussed in the context of the behavioural results, this interaction is the profile that is characteristic for Bayesian Causal Inference: The brain integrates sensory signals at low spatial disparity (i.e. small difference for auditory vs. visual report), but computes different spatial estimates for auditory and visual signals at large spatial disparities (see Fig 4D, Table 2).

**Table 2.** Statistical significance of main, interaction and simple main effects for the behavioural and neural audiovisual weight indices (w_AV_) (‘model free’ approach). TR = task relevance (visual, V or auditory, A, report), VR = visual reliability (high or low), D = disparity (small or large). Asterisks (*) denote significant results (p < 0.05, corrected at the cluster level for multiple comparisons), n.s. = not significant (p ≧ 0.05)

|  | Behavioural | Neural |
| --- | --- | --- |
| VR | $p = 0.61$ | 65 – 510 ms: $p = 0.0002^*$ |
| TR | $p = 0.0002^*$ | 190 – 700 ms: $p = 0.0002^*$ |
| D | $p = 0.77$ | 55 – 130 ms: $p = 0.0062^*$ |
| VR x TR | $p = 0.0002^*$ | n.s. |
| VR x D | $p = 0.0014^*$ | 55 – 135 ms: $p = 0.0002^*$<br>170 – 235 ms: $p = 0.016^*$ |
| TR x D | $p = 0.0002^*$ | 310 – 440 ms: $p = 0.004^*$<br>510 – 590 ms: $p = 0.021^*$ |
| VR x TR x D | $p = 0.79$ | n.s. |
| VR in A | $p = 0.0002^*$ | not tested |
| VR in V | $p = 0.43$ | not tested |
| VR in D - | $p = 0.47$ | 105 – 165 ms: $p = 0.008^*$<br>230 - 515 ms: $p = 0.0002^*$ |
| VR in D + | $p = 0.92$ | 55 – 380 ms: $p = 0.0002^*$ |
| TR in D - | $p = 0.0002^*$ | 230 – 535 ms: $p = 0.0002^*$<br>570 – 630 ms: $p = 0.024^*$ |
| TR in D + | $p = 0.0002^*$ | 235 – 670 ms: $p = 0.0002^*$ |

In addition to these key findings, we also observed a brief but pronounced significant main effect of spatial disparity on wAV at about 55 – 130 ms. While a sound attracted the decoded spatial location at small spatial disparity (i.e. w_AV_ is shifted below 90°, Fig 4 C solid line), the decoded location is shifted away from the sound location (i.e. a repulsive effect) at large spatial disparity (i.e. wAV values above 90°, Fig 4 C, dashed line). Moreover, in this early time window, which coincides with the visual evoked N100 response, the decoded spatial estimate was overall dominated by the visual stimulus location (i.e w_AV_ was close to 90° for both small and large disparity). The effect of disparity may indicate that early multisensory processing is already influenced by a spatial window of integration (Fig 4C, Table 2). Auditory stimuli affected the decoded spatial representations mainly when they were close in space with the visual signal. However, because spatial disparity was inherently correlated with the eccentricity of the audiovisual signals by virtue of our factorial and spatially balanced design, these two effects cannot be fully dissociated. While signals were presented parafoveally or peripherally for small-disparity trials, they were presented always in the periphery for large-disparity trials.

For completeness, we also observed a significant interaction between spatial disparity and visual reliability between 55 – 135 ms and between 170 – 235 ms (Table 2). This interaction results from a larger spatial window of integration for stimuli with low versus high visual reliability. Basically, it is easier to determine whether two signals come from different sources when the visual input is reliable leading to a smaller window of integration.

Finally, we asked whether the neural audiovisual weights were related to the audiovisual weights that observers applied at the behavioural level. Hence, we computed the correlation between the values of the behavioural and neural weight indices w_AV_ separately for each time point. The Fisher z-transformed correlation coefficient fluctuated around chance level until about 100 ms. From 100 ms onwards it progressively increased over time, until it peaked and reached a plateau at about 350 ms (R = 0.72). As expected, this coincides with the time window where we observed a significant interaction between task-relevance and spatial disparity, i.e. the profile characteristic for Bayesian Causal Inference. After 500 ms, it then slowly decreased towards the end of the trial. Cluster permutation test confirmed, that the correlation between neural and behavioural weight indices w_AV_ was significantly better than chance, revealing two significant clusters between 175 – 550 ms (p = 0.0012) and 575 – 665 ms (p = 0.013). These results indicate that the neural representations expressed in EEG activity patterns are critical for guiding observers’ responses.

##### Bayesian modelling analysis

In the EEG Bayesian modelling analysis we fitted five models to the spatial estimates decoded from EEG activity patterns separately for each time point: (i) ‘full segregation audiovisual’, (ii) ‘forced fusion’, (iii) the ‘Bayesian Causal Inference’, (iv) the ‘segregation auditory’, (v) the ‘segregation visual’ models (Fig 3C). The ‘segregation visual’ and ‘segregation auditory’ models incorporate the hypothesis that neural generators may represent only the visual (or only the auditory) location irrespective of whether the visual (resp. auditory) location needs to be reported. In other words, they model a purely unisensory visual (or auditory) source. By contrast, the full segregation audiovisual model embodies the hypothesis that a neural source represents the task-relevant location, i.e. the auditory location for auditory report and the visual location for visual report.

At the random effects group level, Bayesian model comparison revealed a sequential pattern of protected exceedance probabilities across time (Fig 4F): Initially, the ‘segregation visual’ model dominated until about 100 ms post stimulus. This converges with our w_AV_ analysis showing that spatial representations decoded from early EEG activity patterns are dominated by the location of the visual signal (cf. wAV is close to 90°). From 100 to about 200 ms the forced fusion model outperformed the other models indicating that spatial estimates are now influenced jointly by the locations of auditory and visual signals irrespective of their spatial disparity or task-relevance. Again, this mirrors our w_AV_ results where we observed a significant effect of reliability on w_AV_ early (i.e. as expected for forced fusion), while the effect of task-relevance arose later and became prominent from 250 ms onwards.

Hence, both w_AV_ and Bayesian modelling analyses suggest that in this early time window audiovisual signals are predominantly integrated weighted by their reliability into a unified spatial representation irrespective of task-relevance as predicted by forced fusion models. From about 200 ms onwards the protected exceedance probability of the ‘Bayesian Causal Inference’ model progressively increased peaking with an exceedance probability of >0.85 at about 350 ms followed by a plateau until 500 ms. Thus, consistent with the w_AV_ results, audiovisual interactions consistent with Bayesian Causal Inference emerge relatively late at about 350 ms post-stimulus.

## Discussion

Integrating information from vision and audition into a coherent representation of the space around us is critical for effective interactions with the environment. This EEG study temporally resolved the neural dynamics that enables the brain to flexibly integrate auditory and visual signals into spatial representations in line with the predictions of Bayesian Causal Inference.

Auditory and visual senses code spatial location in different reference frames and representational formats [25]. Vision provides spatial information in eye-centred, audition in head-centred reference frames [26,27]. Furthermore, spatial location is directly coded in the retinotopic organization in primary visual cortex [28], while spatial location in audition is computed from sound latency and amplitude differences between the ears starting in the brainstem [26]. In auditory cortices of primates, spatial location is thought to be represented by neuronal populations with broad tuning functions [29,30]. In order to merge spatial information from vision and audition the brain thus needs to establish coordinate mappings and/or transform spatial information into partially shared ‘hybrid’ reference frames as previously suggested by neurophysiological recordings in non-human primates [29,31]. In the first step we therefore investigated the neural dynamics of spatial representations encoded in EEG activity patterns separately for unisensory auditory and visual signals using the method of temporal generalization matrices [21]. In vision, spatial location was encoded initially at 60 ms in transient neural activity associated with the early P1 and N1 components and then turned into temporally more stable representations from 200 ms and particularly from 350 ms (Fig 2, upper right quadrant, Fig S2). In audition, spatial location was encoded by relatively stable EEG activity from 95 ms and particularly from 250 ms, which is associated with the auditory long latency P2 component [22–24] (Fig S3).

Critically, activity patterns encoding spatial location not only generalized across time but also across sensory modalities between 160 and 360 ms. As indicated in Fig 2, SVR models trained on visual evoked responses generalized to auditory evoked responses and vice versa (upper left and lower right quadrant, significant cross-sensory generalization encircled by thick grey line). These results suggest that unisensory auditory and visual spatial locations are initially represented by transient and modality-specific activity patterns. Later, at about 200 ms, they are transformed into temporally more stable representations that may rely on neural sources in fronto-parietal cortices that are at least to some extent shared between auditory and visual modalities [22,32,33].

Next, we asked when and how the human brain combines spatial information from vision and audition into a coherent representation of space. Critically, the brain should integrate sensory signals only when they come from a common event, but segregate signals from independent events [1,2,12]. To investigate how the brain arbitrates between sensory integration and segregation we presented observers with synchronous audiovisual signals that varied in their spatial disparity across trials. On each trial observers reported either the auditory or the visual location. Our results show that a concurrent yet spatially disparate visual signal biased observers’ perceived sound location towards the visual location – a phenomenon coined spatial ventriloquist illusion [17,34]. Consistent with reliability-weighted integration, this audiovisual spatial bias was significantly stronger, when the visual signal was more reliable (Fig 1C left, grey solid vs. dashed lines). Further, observers reported different locations for auditory and visual signals and this difference was even greater for large relative to small spatial disparity trials. This significant interaction between spatial disparity and task-relevance indicates that human observers arbitrate between sensory integration and segregation depending on the probabilities of different causal structures of the world that can be inferred from audiovisual spatial disparity.

Using EEG we then investigated how the brain forms neural spatial representations dynamically post-stimulus. Our analysis of the neural audiovisual weight index w_AV_ shows that the spatial estimates decoded from EEG activity patterns are initially dominated by visual inputs (i.e. w_AV_ close to 90°). This visual dominance is most likely explained by the retinotopic representation of visual space which facilitates EEG decoding of space leading to visual predominance (for further discussion, see methods section). From about 65 ms onwards visual reliability significantly influenced w_AV_ (Fig 4A): As expected the location of the visual signal exerted a stronger influence on the spatial estimate decoded from EEG activity patterns when the visual signal was reliable than unreliable. By contrast, the signal’s task-relevance influenced the audiovisual weight index only later from about 190 ms (Fig 4B). Thus, visual reliability as a bottom-up stimulus-bound factor impacted the sensory weighting in audiovisual integration prior to top-down effects of task-relevance. Most importantly, we observed a significant interaction between task-relevance and spatial disparity as the characteristic profile for Bayesian Causal Inference from about 310 ms: the difference in w_AV_ between auditory and visual report was significantly greater for large than small disparity trials (Fig 4D, Table 2). Thus, spatial disparity determined the influence of task-irrelevant signals on the spatial representations encoded in EEG activity from about 310 ms onwards. A task-irrelevant signal influenced the spatial representations mainly when auditory and visual signals were close in space and hence likely to come from a common event, but it had minimal influence, when they were far apart in space. Collectively, our statistical analysis of the audiovisual weight index revealed a sequential emergence of visual dominance, reliability-weighting (from about 100 ms), effects of task-relevance (from about 200 ms) and finally the interaction between task-relevance and spatial disparity (from about 310 ms, Fig 4A-D).

This multi-stage process was also mirrored in the time course of exceedance probabilities furnished by our formal Bayesian model comparison: The unisensory visual segregation model was the winning model for the first 100 ms - thereby modelling the early visual dominance. The audiovisual forced fusion model embodying reliability-weighted integration dominated the time interval of 100 - 250 ms. Finally, the Bayesian Causal Inference model that enables the arbitration between sensory integration and segregation depending on spatial disparity outperformed all other models from 350 ms onwards. Hence, both our Bayesian modelling analysis and our w_AV_ analysis showed that the hierarchical structure of Bayesian Causal Inference is reflected in the neural dynamics of spatial representations decoded from EEG. Importantly, the Bayesian Causal Inference model also outperformed the audiovisual full segregation model that enables the representation of the location of the task-relevant stimulus unaffected by the location of the task-irrelevant stimulus. Instead, our Bayesian modelling analysis confirmed that from 350 ms onwards the brain integrates audiovisual signals weighted by their bottom-up reliability and top-down task-relevance into spatial priority maps [35,36] that take into account the probabilities of the different causal structures consistent with Bayesian Causal Inference. Critically, the spatial priority maps were behaviourally relevant for guiding spatial orienting and actions as indicated by the correlation between the neural and behavioural audiovisual weight indices, which progressively increased from 100 ms and culminated at about 300-400 ms.

The timing and the parietal-dominant topographies of the AV potentials (see Fig S2, S3) that form the basis for our spatial decoding (and hence for w_AV_ and Bayesian modelling analyses) closely match the P3b component, i.e. a subcomponent of the classical P300. While it is thought that the P3b relies on neural generators located mainly in parietal cortices [37,38], its specific functional role remains controversial [39]. Given its sensitivity to stimulus probability [40–42] and discriminability [43] as well as task-context [39,44,45], it was proposed to reflect neural processes involved in transforming sensory evidence into decisions and actions [46]. Most recent research has suggested that the P3b may sustain processes of evidence accumulation [47] that are influenced by observer’s priors [48], incoming evidence (i.e. likelihood [49]) and observers’ belief updating [50]. Likewise, our supplementary time-frequency analyses revealed that alpha/beta power which has previously been associated with the generation of the P3b component [51] depended on bottom-up visual reliability between 200 – 400 ms and top-down task relevance between 350-550 ms post stimulus (see Fig S5 and Table S2) - thereby mimicking the temporal evolution of bottom-up and top-down influences observed in our main w_AV_ and Bayesian modelling analysis.

Yet, our main analysis took a different approach. Rather than focusing on the effects of visual reliability, task-relevance/attention and spatial disparity directly on ERPs or time-frequency power, the w_AV_ analysis investigated how these manipulations affect the spatial representations encoded in EEG activity patterns and the Bayesian modelling analysis accommodated those effects directly in the computations of Bayesian Causal inference. Along similar lines, two recent fMRI studies characterized the computations involved in integrating audiovisual spatial inputs across the cortical hierarchy [14,16]: While low level auditory and visual areas predominantly encoded the unisensory auditory or visual locations (i.e. full segregation model) [52–61], higher order visual areas and posterior parietal cortices combined audiovisual signals weighted by their sensory reliabilities (i.e. forced fusion model) [62–65]. Only at the top of the hierarchy, in anterior parietal cortices, did the brain integrate sensory signals consistent with Bayesian Causal Inference. Thus, the temporal evolution of Bayesian Causal Inference observed in our current EEG study mirrored its organization across the cortical hierarchy observed in fMRI.

Fusing the results from EEG and fMRI studies (see caveats in methods section) thus suggests that Bayesian causal inference in multisensory perception relies on dynamic encoding of multiple spatial estimates across the cortical hierarchy. During early processing multisensory perception is dominated by full segregation models associated with activity in low level sensory areas. Later audiovisual interactions that are governed by forced fusion principles rely on posterior parietal areas. Finally, Bayesian Causal inference estimates are formed in anterior parietal areas. Yet, while our results suggest that full segregation, forced fusion and Bayesian Causal inference dominate EEG activity patterns at different latencies, they do not imply a strictly feed-forward architecture. Instead, we propose that the brain concurrently accumulates evidence about the different spatial estimates and the underlying causal structure (i.e. common vs. independent sources) most likely via multiple feed-back loops across the cortical hierarchy [18,19]. Critically, only after 350 ms is a final perceptual estimate formed in anterior parietal cortices that takes into account the uncertainty about the world’s causal structure and combines audiovisual signals into spatial priority maps as predicted by Bayesian Causal Inference.

## Methods

### Participants

Sixteen right-handed participants gave informed consent to participate in the experiment. Three of those participants did not complete the entire experiment: two participants were excluded based on eye tracking results from the first day (the inclusion criterion was less than 10% of trials rejected because of eye blinks or saccades, see the Eye movement recording and analysis section for details) and one participant withdrew from the experiment. The remaining thirteen participants (7 females, mean age = 22.1; SD = 3.0) completed the three-day experiment and are thus included in the analysis. All participants had no history of neurological or psychiatric illnesses, normal or corrected-to-normal vision and normal hearing. The study was approved by the research ethics committee of the University of Birmingham and was conducted in accordance with the principles outlined in the Declaration of Helsinki.

### Stimuli

The visual (V) stimulus was a cloud of 20 white dots (diameter = 0.43° visual angle, stimulus duration: 50 ms) sampled from a bivariate Gaussian distribution with vertical standard deviation of 2° and horizontal standard deviation of 2° or 12° visual angle presented on a dark grey background (67% contrast). Participants were told that the 20 dots were generated by one underlying source in the center of the cloud. The visual cloud of dots was presented at one of four possible locations along the azimuth (i.e., -10°, -3.3°, 3.3° or 10°).

The auditory (A) stimulus, was a 50 ms long burst of white-noise with 5 ms on/off ramp. Each auditory stimulus was delivered at 75 dB sound pressure level through one of four pairs of two vertically aligned loudspeakers placed above and below the monitor at four positions along the azimuth (i.e., -10°, -3.3°, 3.3° or 10°). The volumes of the 2 x 4 speakers were carefully calibrated across and within each pair to ensure that participants perceived the sounds as emanating from the horizontal midline of the monitor.

### Experimental design and procedure

In a spatial ventriloquist paradigm, participants were presented with synchronous, spatially congruent or disparate visual and auditory signals (Fig 1A, B). On each trial, visual and auditory locations were independently sampled from four possible locations along the azimuth (i.e., -10°, - 3.3°, 3.3° or 10°) leading to four levels of spatial disparity (i.e., 0°, 6.6°, 13.3° or 20°, i.e. as indicated by the grayscale in Fig 1A). In addition, we manipulated the reliability of the visual signal by setting the horizontal standard deviation of the Gaussian cloud to 2° (high reliability) or 14° (low reliability) visual angle. In an inter-sensory selective-attention paradigm, participants reported either their auditory or visual perceived signal location, and ignored signals in the other modality. For the visual modality, they were asked to determine the location of the center of the visual cloud of dots. Hence, the 4 x 4 x 2 x 2 factorial design manipulated (1) the location of the visual stimulus ({-10°, -3.3°, 3.3°, 10°}, i.e., the mean of the Gaussian) (2) the location of the auditory stimulus ({-10°, -3.3°, 3.3°, 10°}) (3) the reliability of the visual signal ({2°,14°}, STD of the Gaussian) and (4) task-relevance (auditory- / visual-selective report) resulting in 64 conditions (Fig 1A). To characterize the computational principles of multisensory integration, we reorganized these conditions into a two (visual reliability: high vs. low) x two (task-relevance: auditory vs. visual report) x two (spatial disparity: ≤ 6.6° vs. > 6.6°) factorial design for the statistical analysis of the behavioural and EEG data. In addition, we included 4 (locations: 10°, -3.3°, 3.3° or 10°) x 2 (visual reliability: high, low) unisensory visual conditions and 4 (locations: 10°, -3.3°, 3.3° or 10°) unisensory auditory conditions. We did not manipulate auditory reliability, because the reliability of auditory spatial information is anyhow limited. Further, the manipulation of visual reliability is sufficient to determine reliability-weighted integration as a computational principle and arbitrate between the different multisensory integration models (see below for Bayesian modelling).

On each trial, synchronous audiovisual, unisensory visual or unisensory auditory signals were presented for 50 ms, followed by a response cue 1000 ms after stimulus onset (Fig 1B). The response was cued by a central pure tone (1000 Hz) and a blue colour change of the fixation cross presented in synchrony for 100 ms. Participants were instructed to withhold their response and avoid blinking until the presentation of the cue. They fixated a central cross throughout the entire experiment. The next stimulus was presented after variable response interval of 2.6-3.1 s.

Stimuli and conditions were presented in a pseudo-randomized fashion. The stimulus type (bisensory vs. unisensory) and task-relevance (auditory vs. visual) was held constant within a run of 128 trials. This yielded four run types: audiovisual with auditory report, audiovisual with visual report, auditory with auditory report and visual with visual report. The task relevance of the sensory modality in a given run was displayed to the participant at the beginning of the run. Further, across runs we counterbalanced the response hand (i.e. left vs. right hand) to partly dissociate spatial processing from motor responses. The order of the runs was counterbalanced across participants. All conditions within a run were presented an equal number of times. Each participant completed 60 runs leading to 7680 trials in total (3840 auditory and 3840 visual localization tasks, i.e. 96 trials for each of the 76 conditions were included in total; apart from the 4 unisensory auditory conditions that included 192 trials). The runs were performed across three days with 20 runs per day. Each day was started with a brief practice run.

### Experimental set up

Stimuli were presented using Psychtoolbox version 3.0.11 [66] (http://psychtoolbox.org/) under MATLAB R2014a (MathWorks Inc.) on a desktop PC running Windows 7. Visual stimuli were presented via a gamma-corrected 30” LCD monitor with a resolution of 2560 x 1600 pixel at a frame rate of 60 Hz. Auditory stimuli were presented at a sampling rate of 44.1 kHz via 8 external speakers (Multimedia) and an ASUS Xonar DSX sound card. Exact audiovisual onset timing was confirmed by recording visual and auditory signals concurrently with a photo-diode and a microphone. Participants rested their head on a chin rest at a distance of 475 mm from the monitor and at a height that matched participants’ ears to the horizontal midline of the monitor. Participants responded by pressing one of four response buttons on a USB keypad with their index, middle, ring and little finger respectively.

### Eye movement recording and analysis

To address potential concerns that results were confounded by eye movements, we recorded participants’ eye movements. Eye recordings were calibrated in the recommended field of view (32° horizontally and 24° vertically) for the EyeLink 1000 Plus system with the desktop mount at a sampling rate of 2000 Hz. Eye position data were on-line parsed into events (saccade, fixation, eye blink) using the EyeLink 1000 Plus software. The ‘cognitive configuration’ was used for saccade detection (velocity threshold = 30°/sec, acceleration threshold = 8000°/sec^2^, motion threshold = 0.15°) with an additional criterion of radial amplitude larger than 1°. Individual trials were rejected if saccades or eye blinks were detected from -100 to 700 ms post-stimulus.

### Behavioural data analysis

Participants’ stimulus localization accuracy was assessed as the Pearson correlation between their location responses and the true signal source location separately for unisensory auditory, visual high reliability and visual low reliability conditions. To confirm whether localization accuracy in vision exceeded performance in audition in both visual reliabilities, we performed Monte-Carlo permutation tests. Specifically, we entered the subject-specific Fisher z-transformed Pearson correlation differences between vision and audition (i.e. visual – auditory) separately for the two visual reliability levels into a Monte-Carlo permutation test at the group level based on the one-sample t-statistic with 5000 permutations ([67]).

### EEG data acquisition

Continuous EEG signals were recorded from 64 channels using Ag/AgCl active electrodes arranged in 10-20 layout (ActiCap, Brain Products GmbH, Gilching, Germany) at a sampling rate of 1000 Hz, referenced at FCz. Channel impedances were kept below 10kΩ.

### EEG pre-processing

Pre-processing was performed with the FieldTrip toolbox [68] (http://www.fieldtriptoolbox.org/). For the decoding analysis raw data were high pass filtered at 0.1 Hz, re-referenced to average reference, and low pass filtered at 120 Hz. Trials were extracted with 100 ms pre-stimulus and 700 ms post-stimulus period and baseline corrected by subtracting the average value of the interval between -100 – 0 ms from the time course. Trials were then temporally smoothed with a 20 ms moving window and down-sampled to 200 Hz (note, that a 20 ms moving average is comparable to a Finite Impulse Response (FIR) filter with a cut-off frequency of 50 Hz). Trials containing artefacts were rejected based on visual inspection. Furthermore, trials were rejected if they included (i) eye blinks, (ii) saccades, or when (iii) the distance between eye fixation and the central fixation cross exceeded 2 degrees, or (iv) participants responded prior to the response cue or (v) there was no response. For event related potentials (ERPs; Fig S2, S3) the pre-processing was identical to the decoding analysis, except that a 45 Hz low-pass filter was applied without additional temporal smoothing with a temporal moving window. Grand average ERPs were computed by averaging all trials for each condition first within each participant, then across participants.

### EEG multivariate pattern analysis

For the multivariate pattern analyses we computed ERPs by averaging over sets of 8 randomly assigned individual trials from the same condition. To characterize the temporal dynamics of the spatial representations (see below) we trained linear support vector regression models (SVR, LIBSVM [69], https://www.csie.ntu.edu.tw/~cjlin/libsvm/) to learn the mapping from ERP activity patterns of the i. unisensory auditory (for auditory decoding), ii. unisensory visual (for visual decoding) or iii. audiovisual congruent conditions (for audiovisual decoding) to external spatial locations separately for each time point (every 5 ms) over the course of the trial (Fig S2, S3, S4). All SVR models were trained and evaluated in a 12-fold stratified cross-validation (12 ERPs/fold) procedure with default hyper parameters (C = 1, *ε* = 0.001). The specific training and generalization procedures were adjusted to the scientific questions (see below for details)

### Overview of behavioural and EEG analysis

Combining psychophysics, computational modelling and EEG we addressed two questions: First, focusing selectively on the unisensory auditory and unisensory visual conditions we investigated when spatial representations are formed that generalize across auditory and visual modalities. Second, focusing on the audiovisual conditions we investigated when and how human observers integrate audiovisual signals into spatial representations that take into account the observer’s uncertainty about the world’s causal structure consistent with Bayesian Causal Inference. In the following we will describe the analysis approaches to address these two questions in turn.

### Shared and distinct neural representations of space across vision and audition

First, we investigated how the brain forms spatial representations in either audition or vision using the so-called temporal generalization method [21]. Here, the SVR model is trained at time point t to learn the mapping from e.g. unisensory visual (or auditory) ERP pattern to external stimulus location. This learnt mapping is then used to predict spatial locations from unisensory visual (or auditory) ERP activity patterns across all other time points. Training and generalization were applied separately to unisensory auditory and visual ERPs. To match the number of trials for auditory and visual conditions, we applied this analysis to the visual ERPs pooled over the two levels of visual reliability. The decoding accuracy as quantified by the Pearson correlation coefficient between the true and decoded stimulus locations is entered into a training time x generalization time matrix. The generalization ability across time illustrates the similarity of EEG activity patterns relevant for encoding features (i.e. here: spatial location) and has been proposed to assess the stability of neural representations [21]. Basically, if stimulus location is encoded in EEG activity patterns that are stable (or shared) across time, then a SVR model trained at time point t will be able to correctly decode stimulus location from EEG activity patterns at other time points. By contrast, if stimulus location is represented by transient or distinct EEG activity patterns across time, then a SVR model trained at time point t will not be able to decode stimulus location from EEG activity patterns at other time points. Hence, entering Pearson correlation coefficients as a measure for decoding accuracy for all combinations of training and test time into a temporal generalization matrix has been argued to provide insights into the stability of neural representations whereby the spread of significant decoding accuracy to off-diagonal elements of the matrix indicates temporal generalizability or stability [21].

Second, to examine whether and when neural representations are formed that are shared across vision and audition, we generalized not only to ERP activity patterns across time from the same sensory modality, but also from the other sensory modality (i.e. from vision to audition and vice versa). This cross-sensory generalization reveals neural representations that are shared across sensory modalities.

To assess whether decoding accuracies were better than chance we entered the subject-specific matrices of the Fisher z-transformed Pearson correlation coefficients into a between subjects Monte-Carlo permutation test using the one-sample t-statistic with 5000 permutations ([67], as implemented in the FieldTrip toolbox). To correct for multiple comparisons within the two-dimensional (i.e. time x time) data, cluster-level inference was used based on the maximum of the summed t-values within each cluster (‘maxsum’) with a cluster defining threshold of p < 0.05 and a two-tailed p-value was computed.

### Computational principles of audiovisual integration: General linear model-based analysis of audiovisual weight index w_AV_ and Bayesian modelling analysis

To characterize how human observers integrate auditory and visual signals into spatial representations at the behavioural and neural levels, we developed a GLM-based analysis of an audiovisual weight index w_AV_ and a Bayesian Modelling analysis that we applied to both (i) the reported auditory and visual spatial estimates (i.e., participants’ behavioural localization responses) and (ii) the neural spatial estimates decoded from EEG activity pattern evoked by audiovisual stimuli (see Fig 3 and [14,16]).

### General linear model analysis of audiovisual weight index w_AV_

#### Support vector regression to decode spatial estimates from audiovisual EEG activity pattern

The neural spatial estimates were obtained by training a SVR model on the audiovisual congruent trials to learn the mapping from audiovisual ERP activity pattern at time t to external stimulus locations. This learnt mapping at time t was then used to decode the stimulus location from the ERP activity patterns of the spatially congruent and incongruent audiovisual conditions at time t (see Fig 3A and B right). These training and generalization steps were repeated across all times t to obtain distributions of neural (i.e. decoded) spatial estimates for all 64 conditions for every time point t.

#### Regression model and computation of behavioural and neural audiovisual weight index w_AV_

In the ‘GLM-based’ analysis approach we quantified the influence of the location of the auditory and visual stimuli on the reported (behavioural) or decoded (neural) spatial estimates using a linear regression model (see Fig 3C left). In this regression model, the reported (or decoded) spatial locations were predicted by the true auditory and visual stimulus locations for each of the eight conditions in the 2 (visual reliability: high vs. low) x 2 (task-relevance: auditory vs. visual report) x 2 (spatial disparity: ≤ 6.6° vs. > 6.6°) factorial design (Fig 1A).

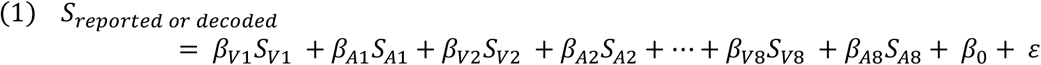

Hence, the regression model included 16 regressors in total, i.e., 8 (conditions) x 2 (true auditory or visual spatial locations). We computed one behavioural regression model for participants’ reported locations. Further, we computed 161 neural regression models for the spatial locations decoded from EEG activity pattern across time, i.e. one neural regression model for every 5 ms interval leading to time courses of auditory (ß_A_) and visual (ß_V_) parameter estimates.

In each regression model the auditory (ß_A_) and visual (ß_V_) parameter estimates quantified the influence of auditory and visual stimulus locations on the reported (or decoded) stimulus location for a particular condition. A positive ß_V_ (or ß_A_) indicates that the true visual (or auditory) location has a positive weight and hence an attractive effect on the reported or decoded location (e.g. it is shifted toward the true visual location, see Fig 3C left for an example). A negative ß_V_ (or ß_A_) indicates that the true visual (or auditory) location has a negative weight and hence a repulsive effect on the reported or decoded location (e.g. it is shifted away from the true visual location). Critically, the auditory and visual parameter estimates need to be interpreted together. To obtain a summary index we computed the relative audiovisual weight (w_AV_) as the four-quadrant inverse tangent of the visual (ß_V_) and auditory (ß_A_) parameter estimates for each of the eight conditions in each regression model (see Fig 3C left). The angles in radians are then converted to degrees:

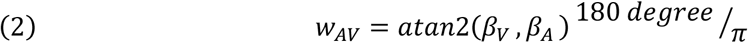

The four-quadrant inverse tangent was used to map each combination of (positive or negative) visual (ß_V_) and auditory (ß_A_) parameter uniquely to a value in the closed interval [-π, π], which was then transformed into degrees. If the reported/decoded estimate is dominated purely and positively by the visual signal (i.e. ß_A_ = 0, ß_V_ > 0), then w_AV_ is 90°. For pure (and positive) auditory dominance, it is 0° (i.e. ß_A_ > 0, ß_V_ = 0). Further, if the visual signal has an attractive influence (i.e. it attracts the perceived location towards the visual location), but the auditory signal has a repulsive influence (i.e. it shifts the perceived location away from the auditory location) on perceived/decoded location (i.e. ß_A_ < 0, ß_V_ > 0), then w_AV_ is > 90° (e.g. Fig. 4 C, high disparity condition).

We obtained one w_AV_ for each of the 8 conditions at the behavioural level and one w_AV_ for each of the 8 conditions and time point (every 5 ms) at the neural level. The neural w_AV_ time courses were temporally smoothed using a 20 ms moving average filter.

#### Statistical analysis of circular indices w_AV_ for behavioural and neural data

We performed the statistics on the behavioural and neural audiovisual weight indices using a two (auditory vs. visual report) x two (high vs. low visual reliability) x two (large vs. small spatial disparity) factorial design based on the likelihood ratio statistics for circular measures (LRTS) [70]. Similar to an analysis of variance for linear data, LRTS computes the difference in log-likelihood functions for the full model that allows differences in the mean locations of circular measures between conditions (i.e. main and interaction effects) and the reduced null model that does not model any mean differences between conditions. LRTS were computed separately for the main effects (i.e. reliability, task-relevance, spatial disparity) and interactions.

To refrain from making any parametric assumptions, we evaluated the main effects of visual reliability, task-relevance, spatial disparity and their interactions in the factorial design using randomization tests (5000 randomizations). To account for the within-subject repeated-measures design at the second random-effects level, randomizations were performed within each participant. For the main effects of visual reliability, task-relevance and spatial disparity, w_AV_ values were permuted within the levels of the non-tested factors. For tests of the two-way interactions (e.g. spatial disparity x task relevance), we permuted the simple main effects of the two factors of interest within the levels of the third factor [71]. For tests of the three-way interaction, values were freely permuted across all conditions [72]. These statistical tests were performed once for behavioural w_AV_ and independently for each time point between 55 – 700 ms (i.e. 130 tests) for neural w_AV_ (see below for multiple comparison correction across time points).

To assess the similarity between behavioural and neural audiovisual weight (w_AV_) indices, we computed the circular correlation coefficient (as implemented in the CircStat toolbox [73]) between the 8 behaivoural (i.e. constant across time) and 8 neural (i.e. variable across time) w_AV_ from our 2 (high vs. low visual reliability) x 2 (auditory vs. visual report) x 2 (large vs small spatial disparity) factorial design separately for each time point.

Unless otherwise stated, results are reported at p < 0.05. To correct for multiple comparisons across time, cluster-level inference was used based on the maximum of the summed LRTS values within each cluster (‘maxsum’) with an uncorrected cluster defining threshold of p < 0.05 (as implemented in the FieldTrip toolbox).

For plotting circular means of w_AV_ (Fig 1C for behavioral w_AV_, Fig 4A-D for neural w_AV_), we computed the means’ confidence intervals (as implemented in the CircStat toolbox [73]).

### Bayesian modelling analysis

#### Description of Bayesian models and decision strategies

Next, we fitted the full-segregation model(s), the forced-fusion model and the Bayesian Causal Inference model to the spatial estimates that were reported by observers (i.e. behavioural response distribution, Fig 3B left) or decoded from ERP activity patterns at time t (i.e. neural spatial estimate distribution, Fig 3B right). Using Bayesian model comparison, we then assessed which of these models is the best explanation for the behavioural or neural spatial estimates.

In the following, we will first describe the Bayesian Causal Inference model from which we will then derive the forced-fusion and full-segregation models as special cases (details can be found in [2,13–15]).

Briefly, the generative model of Bayesian Causal Inference (see Fig 3C right) assumes that common (*C* = 1) or independent (*C* = 2) causes are sampled from a binomial distribution defined by the common cause prior *P_Common_*. For a common source, the ‘true’ location S_AV_ is drawn from the spatial prior distribution N(μ_AV_, σ_P_). For two independent causes, the ‘true’ auditory (S_A_) and visual (S_V_) locations are drawn independently from this spatial prior distribution. For the spatial prior distribution, we assumed a central bias (i.e., μ = 0). We introduced sensory noise by drawing x_A_ and x_V_ independently from normal distributions centred on the true auditory (resp. visual) locations with parameters σ_A_^2^ (resp. σ_V_^2^). Thus, the generative model included the following free parameters: the common source prior p_common_, the spatial prior variance σ_P_^2^, the auditory variance σ_A_^2^ and the two visual variances σ_V_^2^ corresponding to the two visual reliability levels.

The posterior probability of the underlying causal structure can be inferred by combining the common-source prior with the sensory evidence according to Bayes rule:

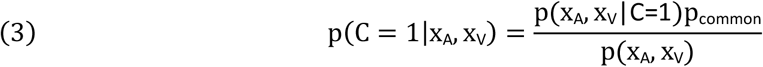

In the case of a common source (C = 1), the optimal estimate of the audiovisual location is a reliability-weighted average of the auditory and visual percepts and the spatial prior (i.e. referred to as forced fusion spatial estimate).

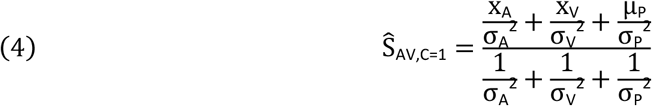

In the case of independent sources (C = 2), the auditory and visual stimulus locations (for the auditory and visual location report, respectively) are estimated independently (i.e. referred to as unisensory auditory or visual segregation estimates):

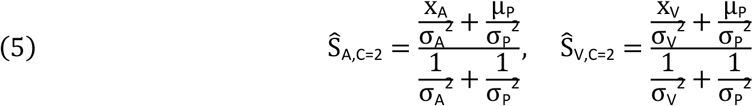

To provide a final estimate of the auditory and visual locations, the brain can combine the estimates from the two causal structures using various decision functions such as ‘model averaging’, ‘model selection’ and ‘probability matching’ [13].

According to the ‘model averaging’ strategy, the brain combines the integrated forced fusion spatial estimate with the segregated, task-relevant unisensory (i.e., either auditory or visual) spatial estimates weighted in proportion to the posterior probability of the underlying causal structures.

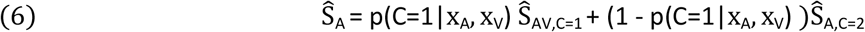

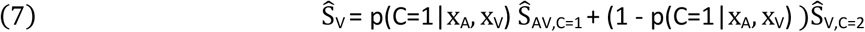

According to the ‘model selection’ strategy, the brain reports the spatial estimate selectively from the more likely causal structure (eq. 8 only shown for Ŝ_A_):

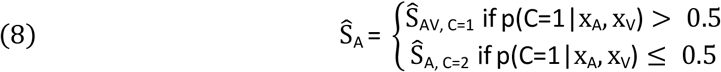

According to ‘probability matching’, the brain reports the spatial estimate of one causal structure stochastically selected in proportion to the posterior probability of this causal structure (eq. 9 only shown for Ŝ_A_):

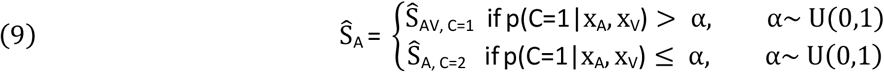

Thus, Bayesian Causal Inference formally requires three spatial estimates (Ŝ_AV,C=ı_, Ŝ_A,C=2_, Ŝ_V, C=2_) which are combined into a final Bayesian Causal Inference estimate (Ŝ_A_ or Ŝ_V_, depending on which sensory modality is task-relevant) according to one of the three decision functions.

In the main paper, we present behavioural results using ‘model averaging’ as the decision function which was associated with the highest model evidence and exceedance probability at the group level. Supplementary Table S1 shows the model evidence, exceedance probabilities and parameters for Bayesian Causal Inference across the three decision strategies for the behavioural data.

At the behavioural level we evaluated whether and how participants integrate auditory and visual stimuli by comparing (i) the Bayesian Causal Inference model (i.e. with model averaging; Table 1), (ii) the forced-fusion model that integrates auditory and visual signals in a mandatory fashion (i.e. formally, the BCI model with a fixed p_common_ = 1, Fig 3C, encircled in yellow) and (iii) the full-segregation model that estimates stimulus location independently for vision and audition (i.e. formally, the BCI model with a fixed p_common_ = 0, i.e. Fig 3C, SegV,A encircled in light blue). This audiovisual full segregation model (SegV,A) assumes that observers report Ŝ_A,C=2_, when they are asked to report the auditory location, and Ŝ_VC=2_, when they are asked to report the visual location. In short, the audiovisual full segregation model (SegV,A) reads out the spatial estimate from the task-relevant unisensory segregation model.

At the neural level, we may also conceive a neural source (or brain region) that represents Ŝ_VC=2_, irrespective of which sensory modality needs to be reported (i.e. Fig 3C, SegV model, encircled in red). For instance, primary visual cortices may be considered predominantly unisensory with selective representations of the visual location even if the observer needs to report the auditory stimulus location. Likewise, we included a model that selectively represents the auditory location (i.e. Fig 3C, SegA model, encircled in green). By contrast, the full segregation audiovisual model (i.e. SegV,A, encircled in light blue) can be thought of as a neural source (or brain area) that encodes the task-relevant estimate computed in a full segregation model. It differs from the Bayesian Causal Inference model by not allowing for any audiovisual interactions or biases irrespective of the probabilities of the world’s causal structure (i.e. operationally manipulated by spatial disparity in the current experiment).

At the behavioural level, the unisensory SegV and SegA models are not useful, because we would expect observers to follow instructions and report their auditory estimate for the auditory report conditions and their visual estimate for the visual report conditions. In other words, it does not seem reasonable to fit the unisensory SegV and SegA models jointly to visual and auditory localization responses at the behavioural level. By contrast, at the neural level, spatial estimates decoded from EEG activity patterns may potentially reflect neural representations that are formed by ‘predominantly unisensory’ neural generators (e.g. primary visual cortex) particularly in early processing phases. Hence, we estimated and compared three models for the behavioural localization reports and five models for the spatial estimates decoded from EEG activity patterns.

#### Model fitting to behavioural and neural spatial estimates and Bayesian Model Comparison

We fitted each model individually to participant’s behavioural localization responses (or spatial estimates decoded from EEG activity pattern at time t) based on the predicted distributions of the spatial estimates (i.e., p(Ŝ|S_A_,S_V_), we use Ŝ as a variable to refer generically to any spatial estimate) for each combination of auditory (S_A_) and visual (S_V_) source location. These predicted distributions marginalize over the internal sensory inputs (i.e. x_A_, x_V_) that are unknown to the experimenter (see [2] for further explanation). More specifically, we fit i. the Bayesian Causal Inference model based on p(Ŝ_A_|S_A_, S_V_) for auditory report conditions and p(Ŝ_V_|S_A_,S_V_) for visual report conditions, ii. the forced fusion model based on p(Ŝ_A V, C=1_|S_A_, S_V_) and iii. the audiovisual full segregation model based on p(Ŝ_A, C=2_|S_A_, S_V_) for auditory report conditions and p(Ŝ_V,C=2_ |Ŝ_A_,S_V_) for visual report conditions. At the neural level, we also fit the unisensory visual segregation model based on p(Ŝ_V,C=2_ |S_A_,S_V_) and the unisensory auditory full segregation model based on p(Ŝ_A,C=2_|S_A_,S_V_) to the spatial estimates decoded from EEG activity pattern across both visual and auditory report conditions.

To marginalize over the internal variables x_A_ and x_V_ that are not accessible to the experimenter the predicted distributions were generated by simulating x_A_ and x_V_ 10000 times for each of the 64 conditions and inferring the different sorts of spatial estimate Ŝ from equations (3)–(9). To link any of those p(Ŝ|S_A_,S_V_) to participants’ auditory and visual discrete localization responses at the behavioural level, we assumed that participants selected the button that is closest to Ŝ and binned the Ŝ accordingly into a histogram (with four bins corresponding to the four buttons). Thus, we obtained a histogram of predicted localization responses for each of those five models separately for each condition and individually for each participant. Based on these histograms we computed the probability of a participant’s counts of localization responses using the multinomial distribution (see [2]). This gives the likelihood of the model given participants’ response data. Assuming independence of conditions, we summed the log likelihoods across conditions.

At the neural level, we first binned the spatial estimates decoded from each ERP activity pattern at each time point based on their distance from the four true locations (i.e., -10°, -3.3°, 3.3° or 10°) into four spatial bins before fitting the models to those discretized spatial estimates.

To obtain maximum likelihood estimates for the parameters of the models (p_common_, σ_P_, σ_A_, σ_V1_ - σ_V2_ for the two levels of visual reliability; formally, the forced fusion and segregation models assume p_common_ = 1 or = 0, respectively), we used a non-linear simplex optimization algorithm as implemented in Matlab’s fminsearch function (Matlab R2016a). This optimization algorithm was initialized with a parameter setting that obtained the highest log likelihood in a prior grid search.

The model fit for behavioural and neural data (i.e. at each time point) was assessed by the coefficient of determination R^2^ [74] defined as

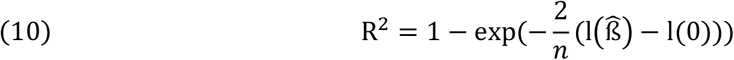

where 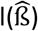 and l(0) denote the log likelihoods of the fitted and the null model, respectively, and n is the number of data points. For the null model, we assumed that an observer randomly chooses one of the four response options, i.e. we assumed a discrete uniform distribution with a probability of 0.25. As in our case the Bayesian Causal Inference model’s responses were discretized to relate them to the four discrete response options, the coefficient of determination was scaled (i.e., divided) by the maximum coefficient (cf. [74]) defined as

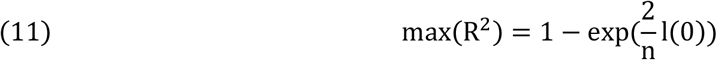

To identify the optimal model for explaining participants’ data (i.e. localization responses at the behavioural level or spatial estimates decoded from EEG activity pattern at the neural level), we compared the candidate models using the Bayesian information criterion (BIC) as an approximation to the model evidence [75].

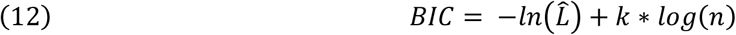

Where 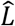 denotes the likelihood, n the number of data points (i.e. EEG activity patterns summed over conditions at a time point t), and k the number of parameters. The BIC depends on both model complexity and model fit. We performed Bayesian model selection [76] at the group (i.e. random effects) level as implemented in SPM8 [77] to obtain the protected exceedance probability that one model is better than any of the other candidate models above and beyond chance.

#### Assumptions and caveats of EEG decoding anlayses

The EEG activity patterns measured across 64 scalp electrodes represent a superposition of activity generated by potentially multiple neural sources located for instance in auditory, visual and higher-order association areas. The extent to which auditory or visual information can be decoded from EEG activity patterns depends therefore inherently on how information is neurally encoded by the ‘neural generators’ in source space and on how these neural activities are expressed and superposed in sensor space (i.e. as measured by scalp electrodes). For example, visual space is retinotopically encoded, while auditory space is represented by broadly tuned neuronal populations (i.e. opponent channel coding model [30,78]), rate-based code [29,79] or spike latency and pattern [80,81]. These differences in encoding of auditory and visual space may contribute to the visual bias we observed for the audiovisual weight index w_AV_ in early processing (Fig 4A-D) and the dominance of the unisensory visual segregation model in the time course of exceedance probabilities (Fig 4F). Further, particularly at later stages scalp EEG patterns likely rely on superposition of activity of multiple neural generators, so that ‘decodability’ will also depend on how source activities combine and project to the scalp (e.g. source orientation etc.). Given the inverse problem involved in inferring sources from EEG topographies recent studies suggested to combine information from fMRI and EEG activity pattern via representational similarity analyses [82,83]. While we informally also pursue this approach in the discussion section of the current paper when merging information from a previous fMRI study that used the same ventriloquist paradigm and analyses with our current EEG results, we recognize the limitations of such a fMRI and EEG fusion approach. For instance, different features encoded in neural activity may be expressed in BOLD-response and EEG scalp topographies [84]. Finally, we trained the support vector regression model on the audiovisual congruent conditions pooled over task-relevance and visual reliability to ensure that the decoder was based on activity patterns generated by sources related to auditory, visual and audiovisual integration processes and the effects of task-relevance or reliability on the audiovisual weight index w_AV_ cannot be attributed to differences in the decoding model (see [62] for a related discussion).

