## Supplementary information for "To integrate or not to integrate: Temporal dynamics of hierarchical Bayesian Causal Inference"

#### Time-Frequency Analysis

##### *Methods*

For time-frequency analysis the pre-processing was identical to the decoding analysis, except that a 100 Hz low-pass filter was applied without additional temporal smoothing and trials were extracted with 400 ms pre-stimulus and 700 ms post-stimulus period time-locked to stimulus onset. EEG data were Fourier transformed with separate parameters for lower (4-30 Hz) and higher (30-80 Hz) frequencies. The time-frequency (TF) powerspectra were computed using sliding time windows of length equal to 3 cycles (low frequencies) or 200 ms (high frequencies) at a given frequency in steps of 2 Hz (low frequencies) or 5 Hz (high frequencies), after application of a Hanning taper (low frequencies) or multitaper with +/- 7 Hz smoothing (high frequencies). The TF powerspectra were averaged across trials within each participant separately for the individual conditions of the two (visual reliability: high vs. low) x two (task-relevance: auditory vs. visual report) x two (spatial disparity: ≤ 6.6° vs. > 6.6°) factorial design. Trials from each condition were randomly sub-selected to ensure an equal number of trials for each of the 8 conditions. TF power was baseline corrected by subtracting the average power within the time window between 400 – 200 ms pre-stimulus from the entire time course separately for each frequency. TF powerspectra were statistically evaluated within the 2x2x2 factorial design using a one-sample t-statistic, computed separately for the main effects (i.e. reliability, task-relevance, spatial disparity) and interactions and separately for high and low frequencies (see above). To refrain from making any parametric assumptions, we used randomization tests (all possible 8192 randomizations). To account for the within-subject repeated-measures design at the second random-effects level, randomizations were performed within each participant ([1], as implemented in the FieldTrip toolbox). To correct for multiple comparisons cluster-level inference was used based on the maximum of the summed t-values within each cluster ('maxsum') with a cluster defining threshold of $p < 0.05$ uncorrected. A two-tailed p-value was computed corrected for multiple comparisons within the time window from 200 ms pre-stimulus to 700 ms post-stimulus x topography (2 dimensional) x frequencies separately for lower (4-30 Hz) and higher (30-80 Hz) frequencies.

##### *Results*

Using one-sample t-tests we assessed the main effects (i.e. reliability, task-relevance, spatial disparity) and interactions separately for high (30-80 Hz) and low (4-30 Hz) frequencies. For the low frequencies, we observed significant effects with time x topography (2 dimensional) x frequency clusters encompassing the alpha/beta-bands (i.e. 8-30 Hz). To characterize these significant clusters that live in a four-dimensional time x topography (2 dimension) x frequency space, we averaged the power within the alpha/beta band (i.e. 8-30 Hz). In Fig S5 we then show the time course of alpha/beta power (A) and the associated topography (B). Please note that we averaged power in the alpha/beta frequency band only for illustrational

purposes, the statistics were performed and corrected for multiple comparisons across the four dimensional time x topography x frequency space.
Generally, audiovisual stimulation induced a transient increase in power relative to baseline in alpha/beta frequency band (50-100 ms post-stimulus), followed by a significant sustained suppression (event related desynchronization, ERD, see Fig S5A and Table S2, i.e. 'overall effect') from 100-700 ms post-stimulus. Visual reliability significantly modulated alpha/beta TF power: Audiovisual stimuli elicited greater ERD when the visual signal was of low than high reliability. The effect was most prominent between 200 – 400 ms post stimulus, predominantly in the alpha/beta frequency band (8-30 Hz) and widespread across the entire scalp (see Fig S5 first row and Table S2). We also observed a significant effect of task relevance both in the low and high frequency bands. In the alpha/beta band, the effect was most prominent between 200 – 0 ms pre-stimulus and 350 – 550 ms post-stimulus and widespread across the scalp (see Fig S5 second row and Table S2). The effect in the high frequencies (30-80 Hz) was observed mostly over fronto-central electrodes between -200 and 700 ms (Table S2). Because the increase in broadband gamma power for auditory relative to visual report could be caused by an increase in the rate of spontaneous miniature saccades [2] that were not excluded during the artefact rejection, we will not interpret this result.
Furthermore, we observed a significant interaction between visual reliability and task relevance in the alpha/beta band over frontal and parieto-occipital channels starting at about 400 ms post stimulus (see Fig S5 fourth row and Table S2). Neither spatial disparity (see Fig S5 third row) nor any of the other two- or three-way interactions were significant.

To summarize, bottom-up visual reliability modulated alpha/beta power between 200-400 ms post-stimulus, while top-down task-relevance influenced alpha/beta power prior to stimulus onset (-200 – 0 ms) and post stimulus (350-550 ms). The pre-stimulus effects of task-relevance could reflect the build-up of expectations and allocation of attentional resources as the task relevance factor was kept constant within each block.

### Supplementary figures

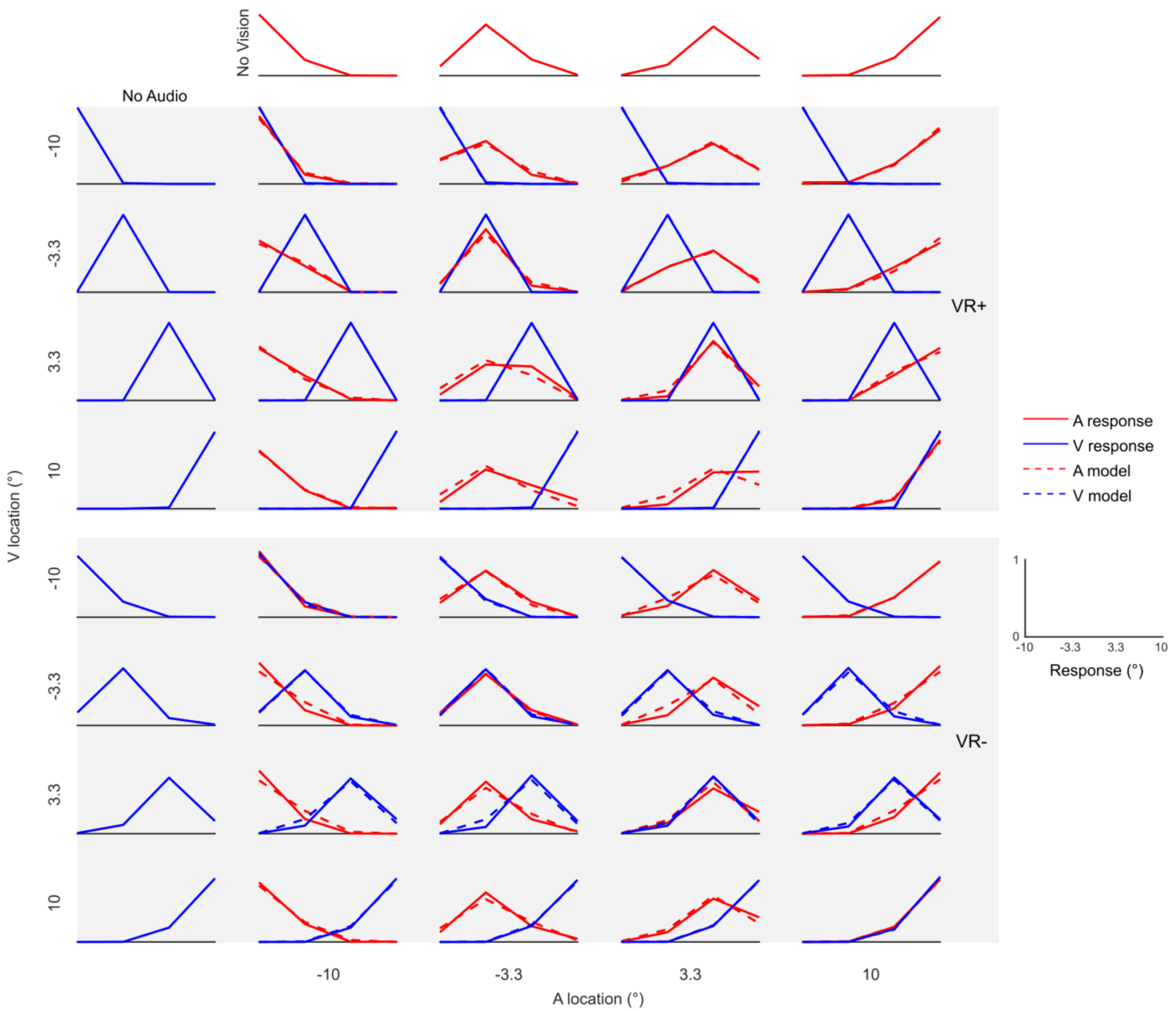


**Figure S1 Distributions of spatial estimates**. The distribution (across participants' mean) of spatial estimates given by observers' behavioural localization responses (solid lines) or predicted by the Bayesian Causal Inference model fitted to observers' behavioural responses (dashed lines, for model averaging) are shown across all conditions in our 2 task-relevance (auditory: red vs. visual: blue) x visual reliability (high: row 1-4 vs. low: row 5-8) x 4 auditory location (columns as indicated) x 4 visual location (rows as indicated) design.

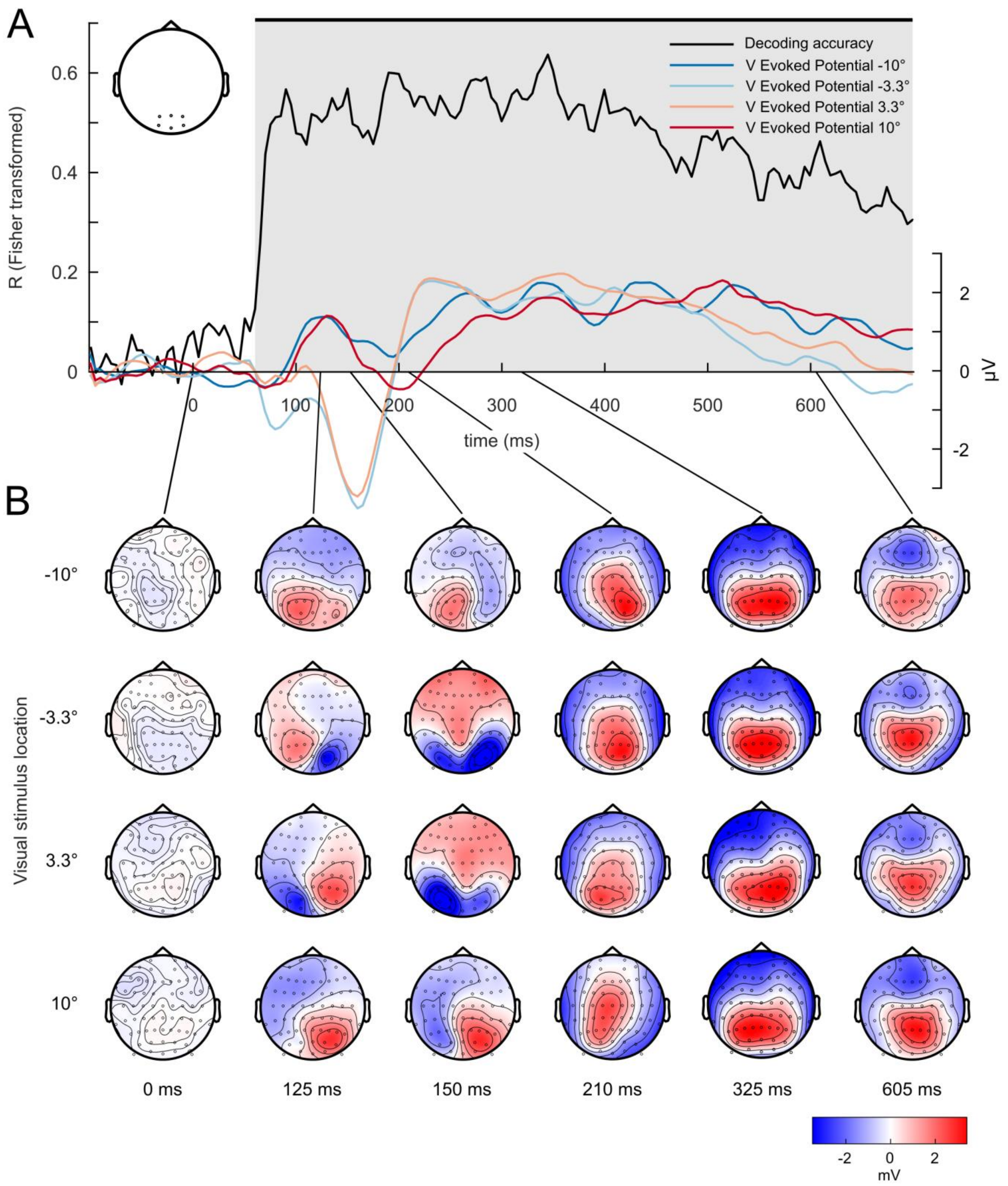


**Figure S2 Time resolved decoding of visual location for unisensory visual stimuli. (A)** Time course of decoding accuracy (i.e. Pearson correlation between true and predicted visual stimulus locations pooled over both visual reliabilities, black line) and the EEG evoked potentials (across participants' mean) for the unisensory visual (high reliability only) signals at -10°, -3.3°, 3.3°, 10° degrees, averaged over occipital channels. Shaded grey area indicates the time window where the decoding accuracy is significantly better than chance. EEG signals were averaged across the electrodes shown in the inset. **(B)** EEG topographies (across participants' mean) for the unisensory visual signals (high reliability only) at -10°, -3.3°, 3.3°, 10° degree shown at the given time points.

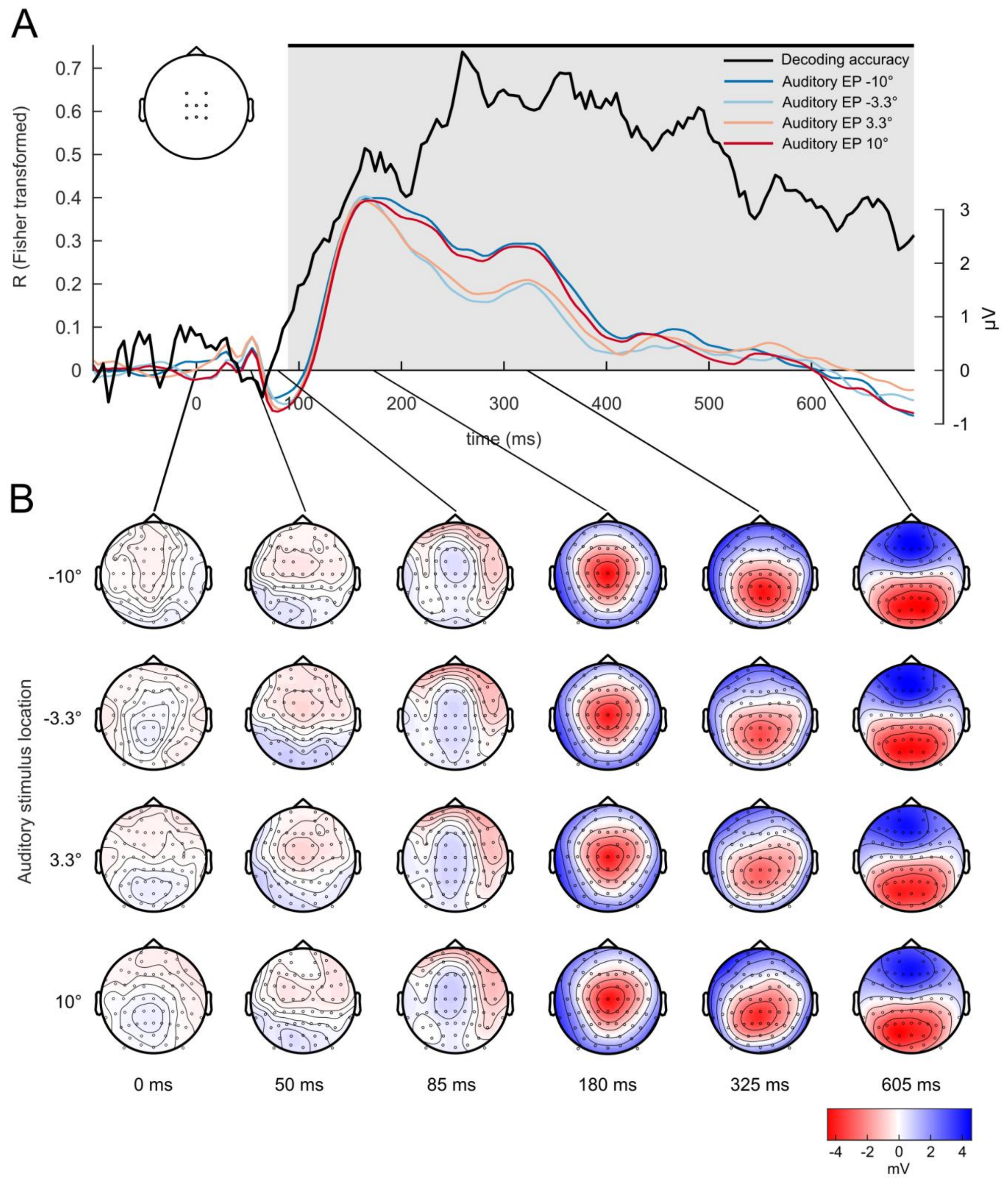


**Figure S3 Time resolved decoding of auditory location for unisensory auditory stimuli. (A)** Time course of decoding accuracy (i.e. Pearson correlation between true and predicted stimulus locations, black line) and the EEG evoked potentials (across participants' mean) for the unisensory auditory signals at -10°, -3.3°, +3.3°, 10° degree, averaged over central channels. Shaded grey area indicates decoding accuracy significantly better than chance. EEG signals were averaged across the electrodes shown in the inset. **(B)** EEG topographies (across participants' mean) for the unisensory auditory stimuli at -10°, -3.3°, 3.3°, 10° degree shown at the given time points.

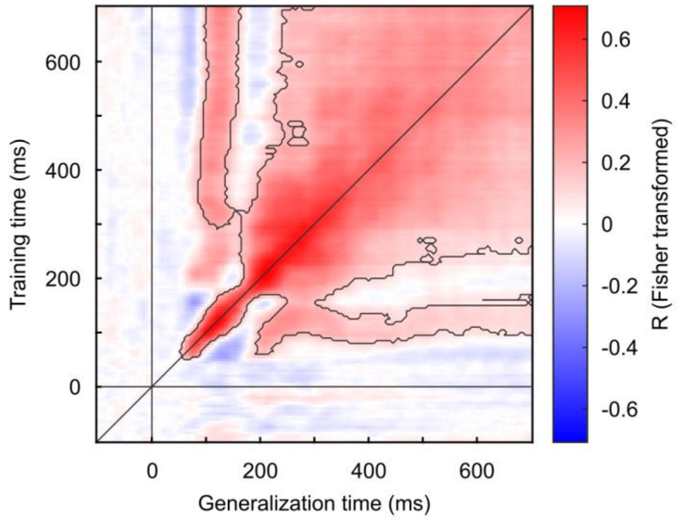


**Figure S4 Temporal generalization matrix for audiovisual congruent trials.** The temporal generalization matrix shows the decoding accuracy for audiovisual congruent trials across each combination of training (y axis) and testing (x axis) time point. The grey line along the diagonal indicates where the training time is equal to the testing time. Horizontal and vertical grey lines indicate the stimulus onset. The thin black lines encircle the cluster with decoding accuracies that were significantly better than chance at $p < 0.05$ corrected for multiple comparisons.

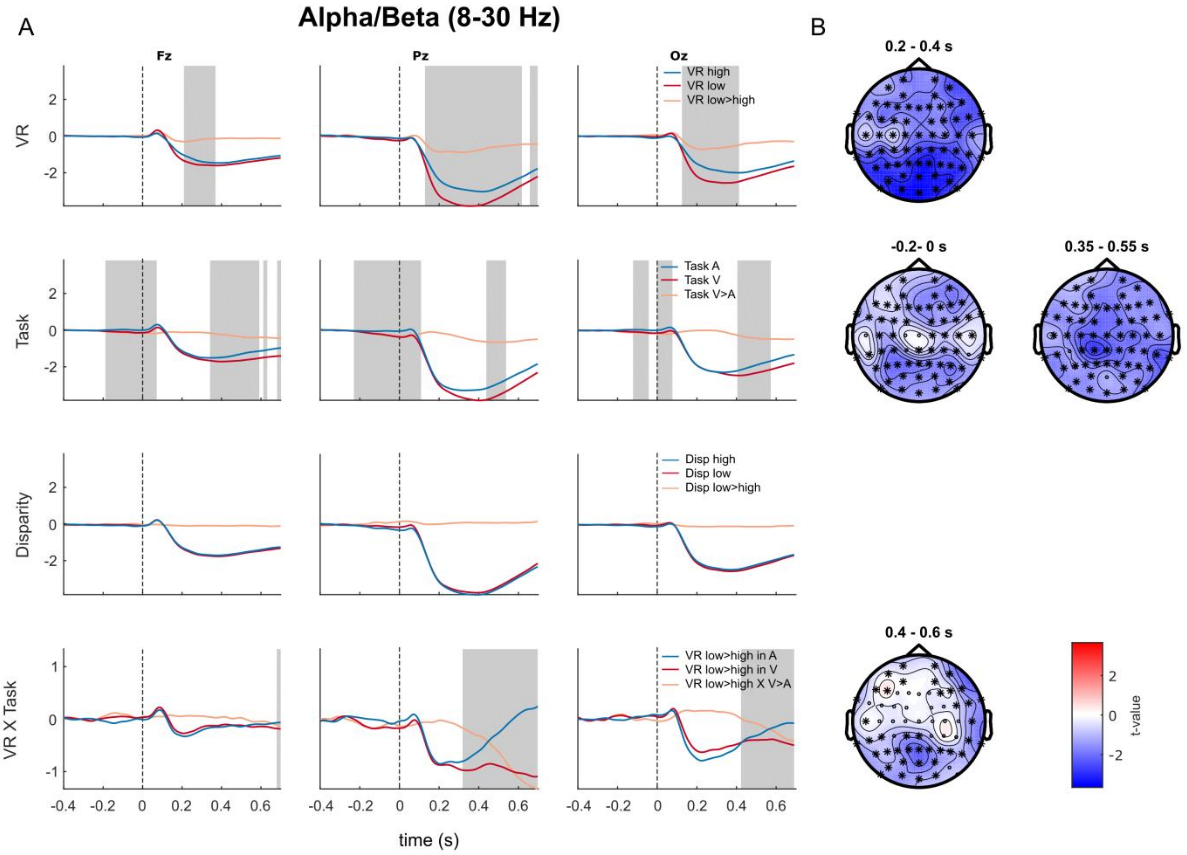


**Figure S5 Time-frequency results for oscillatory power in the alpha / beta band. (A)**. Time courses of total power averaged over the alpha/beta (8-30 Hz) frequency bands (baseline corrected using pre-stimulus window [-400ms - -200ms]) are shown for the main effects of visual reliability (row 1), task relevance (row 2), spatial disparity (row 3) and the visual reliability X task relevance interaction (row 4) at three selected electrodes (i.e. Fz = left; Pz = middle; Oz = right columns). For each effect, we show the power for the difference (or interaction) and the individual conditions coded in different colours as indicated for each row. Grey shaded areas indicate the time windows where at least one electrode was part of the significant cluster after correcting for multiple comparisons across time (i.e. -200 ms to 700 ms), frequency (i.e. 4-30 Hz) and topography. **(B)** Topographies of the t-values averaged across the significant time windows of the corresponding effects. Electrodes marked with black stars were part of the significant cluster (corrected across topography x time x frequency).

### Supplementary tables

| | $p_c$ | $\sigma_p$ | $\sigma_A$ | $\sigma_{V1}$ | $\sigma_{V2}$ | $R^2$ | $relBIC_{group}$ | PEP |
|---|---|---|---|---|---|---|---|---|
| $BCI_{avg}$ | 0.15 ± 0.04 | 36.4 ± 11.0 | 4.4 ± 0.2 | 0.3 ± 0.15 | 3.5 ± 0.24 | 0.857 ± 0.003 | 0 | 0.784 |
| $BCI_{sel}$ | 0.28 ± 0.04 | 25.8 ± 5.3 | 4.4 ± 0.2 | 0.3 ± 0.16 | 3.5 ± 0.23 | 0.856 ± 0.003 | -60.5 | 0.113 |
| $BCI_{match}$ | 0.15 ± 0.03 | 33.0 ± 9.4 | 4.1 ± 0.2 | 0.3 ± 0.16 | 3.4 ± 0.23 | 0.856 ± 0.003 | -54.8 | 0.103 |

**Table S1** Model parameters (across-subjects mean ± SEM) and fit indices of the Bayesian Causal Inference models with different decision functions: model averaging ($BCI_{avg}$), model selection ($BCI_{sel}$) and probability matching ($BCI_{match}$). $R^2$ = coefficient of determination, $relBIC_{group}$ = group level relative BIC, PEP = protected exceedance probability [3]

| | Neural latency | | | | |
|---|---|---|---|---|---|
| Effect | ~ -200 ms | ~ 50 ms | ~ 100 ms | ~ 200 ms | ~ 400 ms |
| Overall | | α/β 50 – 100 ms (p = 0.054) | α/β 100 - 700 ms (p = 0.0001) | | |
| VR | | | | α/β 200 - 400 ms (p = 0.0001) | |
| Task | α/β -200 – 0 ms (p = 0.013) | | | α/β 350 – 550 ms (p = 0.013) | |
| | γ -200 – 700 ms (p = 0.0002) | | | | |
| VR X Disp | | | | | α/β 400 - 700 ms (p = 0.005) |

**Table S2** Time-frequency results. Significant effects are shown for overall relative to baseline, main effect of visual reliability (VR), main effect of task and the interaction between VR and task are shown across rows. Columns of the table indicate the approximate time windows that the significant cluster spanned. All p-values are reported at the cluster level, corrected for multiple comparisons over time x topography x frequency.
